# A Multispecies, Modality-Agnostic Scalable In Vivo Mosaic Screening Platform for Therapeutic Target Discovery

**DOI:** 10.64898/2026.02.26.708253

**Authors:** Vishwaraj Sontake, Vinay Kartha, Neety Sahu, Daniel R. Fuentes, Linda Chio, Hikaru Miyazaki, Jingshu Chen, Arnav Gupta, Jesi Nonora, Arthi Vaidyanathan, Smitha Shambhu, Gayathri Donepudi, Carmen Le, Lianna Fung, Amber Lim, Chase Bowman, Diego Garcia, Dimitry Popov, Kelly Fagan, Charlie Longtine, Juana M. Cruz Sampedro, Samuel B. Hayward, Adam Biedrzycki, Barclay B. Powell, Katie Doroschak, Rachael Watson Levings, Chris Carrico, Ian Driver, Chris Towne, Francisco LePort, Martin Borch Jensen

**Affiliations:** Gordian Biotechnology, South San Francisco, CA USA; Department of Orthopaedic Surgery and Sports Medicine, University of Florida; Department of Large Animal Clinical Sciences, UF

## Abstract

Validating therapeutic targets for complex diseases requires investigating gene functions within native tissue architectures rather than reductionist in vitro models. Here we present a modality-agnostic AAV-based in vivo high-throughput screening platform capable of delivering knockouts, gain-of-function, and synthetic miRNA knockdowns directly to cells within the diseased environment. This system scales to hundreds of perturbations and is adaptable to diverse species and organ systems. To translate high-dimensional screen data into therapeutic assessment, we established a curated analysis framework that scores single-cell transcriptomes against human disease-specific molecular signatures. This method enables quantitative ranking of targets across distinct biological domains ranging from structural fibrosis to inflammatory signaling, to narrow in on the therapeutic potential of each intervention. We applied this strategy to screen loss- and gain-of-function libraries in a murine pulmonary fibrosis model and within the spontaneously osteoarthritic joints of aged horses, identifying metabolic, antifibrotic, and immunomodulatory targets. Importantly, our analysis framework successfully predicted functional outcomes in orthogonal human ex vivo tissue models, including soluble collagen reduction in lung slices and glycosaminoglycan restoration in cartilage, thus establishing a powerful paradigm for prioritizing therapeutic targets by uniting human disease signatures with highly multiplexed in vivo functional genomics.

## Introduction

Clinical translation of novel therapeutic targets is bottlenecked by the lack of scalable validation platforms within predictive preclinical models. While high-throughput CRISPR screens have revolutionized functional genomics in 2D cell cultures^1,2^, these simplified systems often fail to recapitulate spatiotemporal interplay of the complex in vivo microenvironment, including matrix mechanics, vascular and lymphatic gradients, immune–stromal reciprocity, and inflammatory niches, thus limiting their translational value to physiological relevance^3^.

Recent advances in in vivo pooled perturbation screens combined with single-cell RNA sequencing (Perturb-seq) have begun to address this gap^4^. Studies have successfully utilized lentiviral and adeno-associated virus (AAV) vectors to dissect gene function in mouse brain, identifying regulators of neurodevelopmental disorders and glioblastoma within the native tissue^4–7^. Similarly, these methods have been adapted to uncover regulators of metabolism and regeneration in the liver^8^. Despite these advances, the application of high-throughput in vivo perturb-seq to test therapeutic potential at scale remains constrained by three key hurdles. First, the vast majority of studies rely on Cas9-expressing transgenic murine models, precluding genetically diverse models or the use of large animal models that more faithfully recapitulate human disease physiology. Second, current platforms operate predominantly in a single perturbation modality (knockout [*KO*] via transgenic CRISPR–Cas9), limiting our ability to evaluate therapeutic overexpression (*OE*) strategies. Third, while in vivo screens generate vast datasets, their analysis is typically restricted to pathway enrichment and biological exploration. Shifting from standard pathway enrichment to a patient-centric framework, one that evaluates perturbation effects against patient-specific molecular signatures is essential for ranking and prioritizing translationally relevant targets.

To bridge the gap between in vivo discovery and human therapeutic translation, we present an AAV-based high-throughput in vivo screening platform focused on assessing potential therapeutic effects of perturbations. We call this mosaic screening, in reference to the intact organ structure wherein a mosaic of perturbations are evaluated. The workflow integrates target selection, pooled AAV library production, and in vivo incubation, followed by single-cell RNA sequencing (scRNA-seq) for context-specific probing of transcriptomic effects, as for Perturb-seq. We demonstrate that AAV-based mosaic screening works for both *KO* (CRISPR-based dual gRNA constructs), gain-of-function (*GOF*) via *OE* of coding sequences, and *KD (*synthetic miRNA) screens. Further, by combining these perturbations with an analytical framework to map transcriptomic states to human disease signatures, we ensure that identified hits are relevant to human pathophysiology. We validated the predictive power of this framework by demonstrating the functional concordance between perturbation effects and physiological outcome using human primary ex vivo cultures (**Fig 1**).

**Figure 1.**
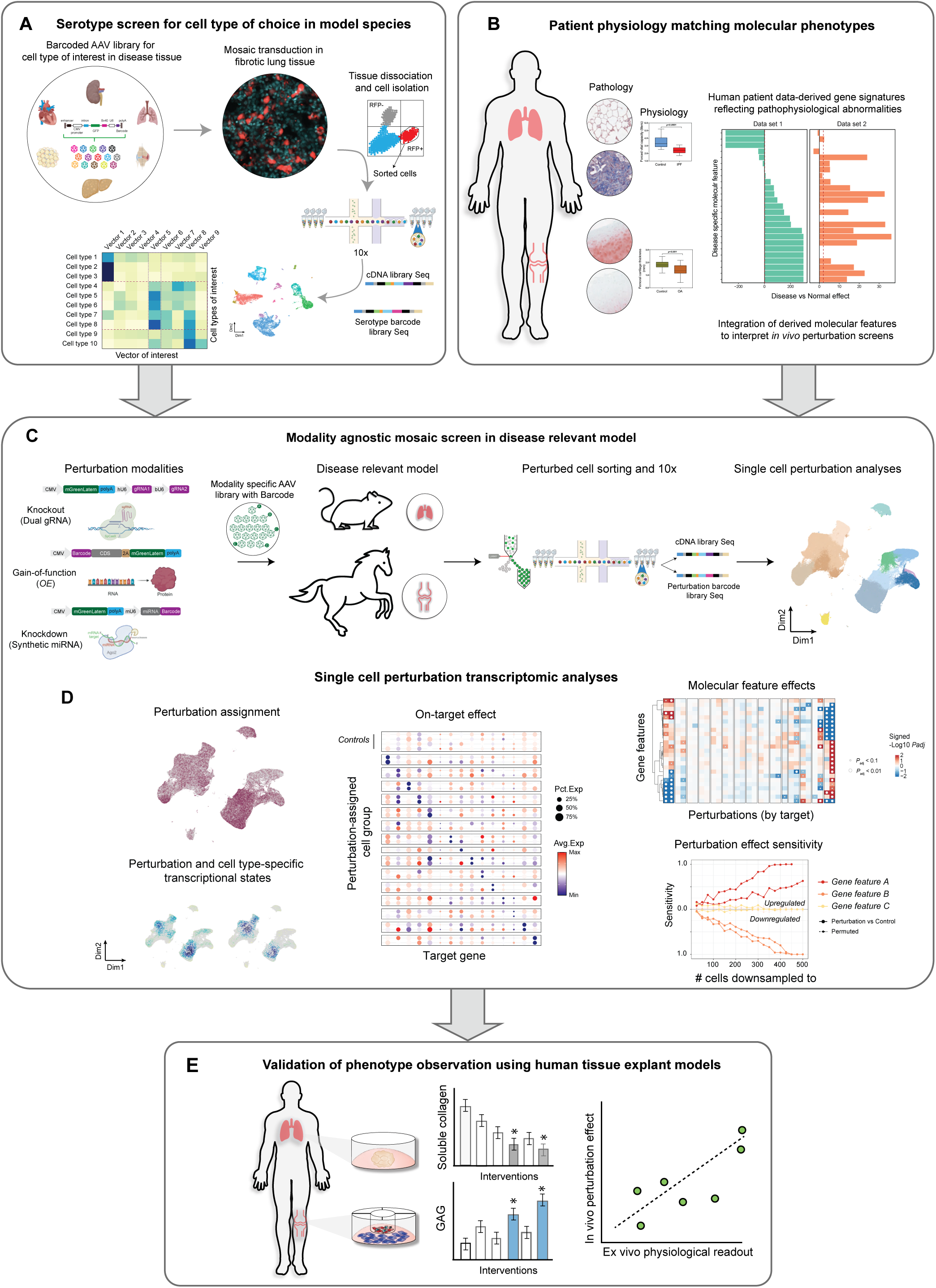
Scalable in vivo mosaic screening platform for target discovery and validation. **A.** schematic depicting AAV serotype screening workflow in disease tissue to optimize specific transduction of target cell type(s). **B.** Paired datasets enable curation of molecular features capturing human pathophysiology to interpret in vivo perturbation data. **C.** Schema showing the modality-agnostic mosaic screening workflow in disease-relevant models**. D.** Single cell computational analysis framework to assess perturbation effects and therapeutic potential using molecular features. **E.** Validation of in vivo screen phenotypes using human primary tissue derived ex vivo models with physiological readouts.

We applied this platform to two distinct, complex diseases of aging: idiopathic pulmonary fibrosis (IPF) and osteoarthritis (OA). Using a progressive pulmonary fibrosis mouse model^9^ we validated distinct roles for *Jak1* and *Tgfbr2* as critical drivers of inflammation and fibrotic states. Through *OE* screens in the same model, we identified *Klf15* as a metabolic regulator of alveolar repair. Extending the platform to a clinically translatable equine model of naturally occurring OA, we successfully identified the *SOCS* family as potent therapeutic targets for synovial inflammation. Crucially, the high-resolution readout also unmasked pleiotropic liabilities of *IL13*, demonstrating that our framework can simultaneously identify mechanistic efficacy and potential therapeutic risks in large-animal models.

## Results

### Mosaic labeling, gRNA capture, and on-target effect validates in vivo CRISPR editing in fibrotic tissue

In vivo perturbation screening in disease-relevant models requires effective delivery of perturbations across a range of pathophysiological states. Fibrosis is a primary driver of tissue dysfunction in age-related diseases, including IPF. To overcome the delivery barriers presented by stiff, remodeled fibrotic tissue in mice, we performed an in vivo screen of 16 barcoded AAV serotypes (see methods) during the active phase of bleomycin-induced injury^10,11^ (**Fig S1A**). Single-cell transcriptomic analysis of the recovered GFP+ fraction captured all major lineages, including injury-specific KRT8+ transitional alveolar type 2 (AT2) cells^12,13^ (**Fig S1B-C**). Barcode enrichment across cell types revealed distinct tropism: while commonly used serotypes (e.g., AAV8, AAV9) showed limited penetrance, AAV6 and AAV-DJ9 emerged as superior vectors with complementary profiles.

AAV6 preferentially targeted epithelial lineages (AT1, AT2, Ciliated), whereas AAV-DJ9 transduced across stromal niche including, matrix producing adventitial fibroblasts, myofibroblasts, pericytes, alveolar fibroblasts, and disease-specific KRT8+ transitional-AT2 cells (**Fig S1D**). This establishes AAV-DJ9 as a potent tool for delivering genetic perturbations to the primary cellular effectors of lung fibrosis in vivo.

To validate our ability to induce and interpret genetic perturbations in vivo, we performed a proof-of-concept CRISPR-Cas9 *KO* screen targeting 15 genes previously implicated in IPF, using a mouse model of progressive pulmonary fibrosis (**Fig2A**). These genes comprised four functional categories: secreted factors (*Thbs1, Postn, Bgn, Fbln1, Cthrc1, Anxa1*), receptors (*Tgfbr2, Lpar1*), kinases (*Pik3ca, Mapk1, Jak1*), and transcriptional regulators (*Atf3, Cebpb, Ctnnb1, Cdkn1a*)^14–17^. To maximize editing efficiency, we designed AAV constructs expressing two gRNAs per target gene (total, 4 gRNAs per gene), with non-targeting and safe-targeting gRNAs as controls^18,19^ (**Fig2A**). The library was administered four weeks after the initial bleomycin administration, once fibrosis was established, with viral dosing optimized for low transduction (<10%) to minimize multiple perturbations per cell and preserve the native pathophysiological environment (**Fig 2B**).

**Figure 2.**
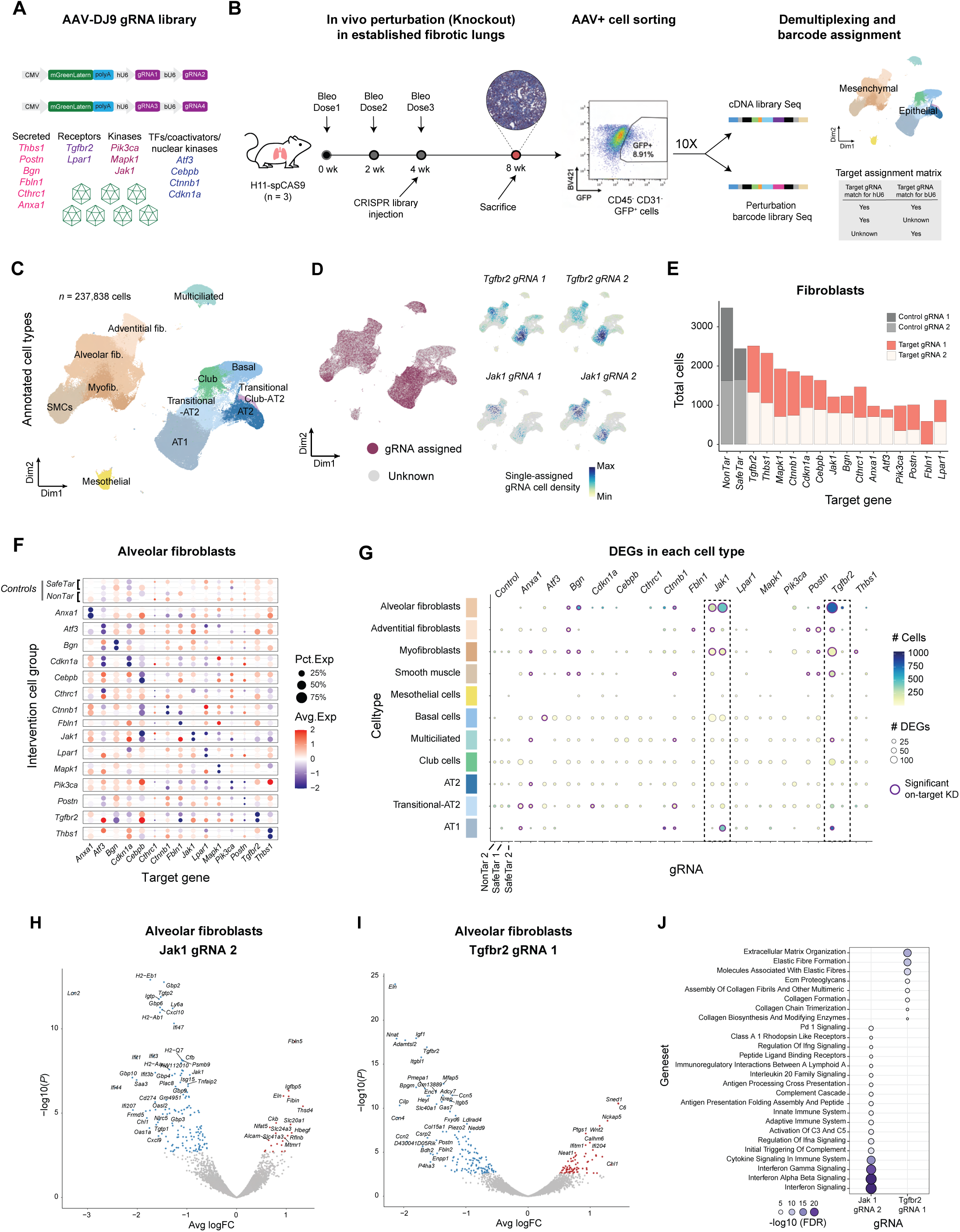
In vivo CRISPR *KO* mosaic screen validated distinct roles *of Jak1 and Tgfbr2* in lung fibrosis. **A.** Schematic depicting dual-gRNA construct design and target classes used in CRISPR screen in PF model. **B.** Schematic depicting the experimental workflow of mosaic screening in PF, including model development, pooled gRNA library delivery, FACS, scRNA-seq data demultiplexing and barcode assignment. **C.** UMAP of cells by annotated cell type indicating mesenchymal and epithelial cell subtypes. **D.** Distribution of gRNA assigned (purple) and non-assigned (gray) cells (left), with density plots showing cell-specific assignment of *Tgfbr2* and *Jak1* gRNAs (right). **E.** Number of single gRNA-assigned fibroblast cells recovered per target genes and control. **F.** Dot plot showing scaled expression of library target genes in alveolar fibroblasts single-assigned to targeting gRNAs or controls (NonTar/SafeTar). Dot size indicates the percentage of cells with detectable gene expression. Color scale indicates Z-score scaled average gene expression across assigned cell groups. **G.** Dot plot showing quantification of differentially expressed genes (DEGs) in each cell type in gRNA assigned cells relative to non-targeting control guide cells (Adj *P < 0.1*). On-target (purple circle) indicates genes where the target gene is downregulated relative to *NonTar* gRNA control cells. **H.** Volcano plots highlighting top differentially expressed genes by *Jak1,* and **I**. *Tgfbr2 LOF* (Adj *P* < 0.1) in alveolar fibroblasts. **J.** Gene set enrichment analysis for significantly down-regulated genes in response to *Jak1* and *Tgfbr2 LOF (hyperenrichment test FDR < 0.01*) in alveolar fibroblasts run using Reactome genesets.

Following a four-week incubation, we sorted CD45⁻/CD31⁻/GFP⁺ cells for scRNA-seq with direct gRNA capture, recovering a total of 237,838 cells across major fibroblast and epithelial lineages (**Fig 2B-C**). Cell identities were confirmed via canonical markers, including *Pdgfra/Tcf21* (alveolar fibroblasts), *Cthrc1/Acta2* (myofibroblasts), and *Krt8/Cldn4* (transitional-AT2) (**Fig S2A-B**). We assigned gRNAs to cells by modeling the observed gRNA unique molecular identifier (UMI) detection rate relative to the expected background rate given droplets comprising ambient RNA (see Methods) (**Fig2D, Fig S2C**), further distinguishing cells receiving single (*n* = 62,828 cells or 26.42% total) versus multiple guides (*n* = 47,072 or 19.79% total) (**Fig S2C-D**). The outer range of cell assignment counts was 68 and 1860 cells per gRNA construct across both fibroblast and epithelial clusters; in fibroblasts most gRNAs were single-assigned to 200-1000 cells, while in epithelial cells most gRNAs were single-assigned to 100-250 (**Fig2E, Fig S2E**). We confirmed gRNA functionality via on-target *KD* efficiency across cell types, observing a significant reduction in target gene expression in alveolar fibroblasts and transitional epithelial cells (**Fig2F and Fig S2F**). Downsampling analysis further demonstrated that 10/15 (∼67%) targets had at least one gRNA that achieved a true positive rate (TPR) of 80% or more for detecting significant knockdown with as few as 200 cells (Wilcoxon DE *P* < 0.05, see Methods) confirming the screen’s sensitivity in detecting gene perturbation within living, fibrotic tissue (**Fig S2G-H**).

### In Vivo CRISPR screen validates *Tgfbr2* and *Jak1* as key fibrotic and inflammatory modulators

Having validated functional perturbation in fibrotic lung cells, we performed differential expression analysis to assess the resulting transcriptome-wide changes across different cell types. Differential gene expression for each perturbation against non-targeting gRNA control cells per cell type suggested cell type-specific effects, with guides targeting *Jak1* and *Tgfbr2* eliciting the largest effects overall (**Fig 2G**). Specially, *Jak1 KO* induced a profound transcriptomic shift in alveolar fibroblasts and myofibroblasts (**Fig 2H**). We observed a broad suppression of immune modules including interferon signaling (*Ifit1, Isg15*), MHC class II antigen presentation (*H2-Ab1, Nlrc5*), and host defense GTPases (*Gbp2, Igtp*), concomitant with the upregulation of structural extracellular matrix (ECM) components (*Fbln5, Eln, Alcam, Thsd4*) (**Fig 2J, left**). This distinct signature suggests that *Jak1* loss effectively decouples the inflammatory and fibrogenic functions of the fibroblast, repressing immune surveillance while promoting matrix organization.

In contrast, *Tgfbr2* KO in alveolar fibroblasts significantly modulated structural programs, downregulating key ECM organizers (*Eln, Postn, Fbln2*) and cytoskeletal/integrin regulators (*Itgbl1, Itgb5, Piezo2*) (Fig 2I). Interestingly, this loss of fibrogenic signaling triggered a compensatory interferon response (*Ifitm1, Ifi204*), reflecting the opposite of the *Jak1* phenotype (**Fig 2I-J**). Pathway enrichment analysis confirmed this dichotomy, i.e., *Jak1* inhibition is predominantly anti-inflammatory, while loss of *Tgfbr2* is fundamentally anti-fibrotic (**Fig 2J**). This functional diversion aligns with the established roles of the TGFβ pathway as a master regulator of fibrosis and *JAK1* as a critical immunomodulator ^20^ ^21–24^, directly reflecting distinct therapeutic mechanisms currently being investigated for their clinical efficacy in IPF patients ^25,26^. Collectively, these findings confirm the ability of our pooled in vivo CRISPR *KO* screen to capture pathway-specific cellular responses within animal models of chronic disease.

### In Vivo *gain of function* screen reveals *Klf15* and *Pparg* as metabolic regulators of lung fibrosis resolution

Gain-of-function (*GOF*) screens are a powerful tool for identifying candidate therapies that require gene activation^27^. While CRISPR activation (CRISPRa) offers context-dependent gene induction, its application in human therapeutics is limited by delivery challenges and chromatin inaccessibility, where tightly packed DNA prevents the CRISPR machinery from binding^28,29^, and off-target ‘neighbor’ activation^30–32^. In contrast, AAV-based *OE* approaches overcome these limitations by decoupling expression from the endogenous chromatin environment, and have already been approved for human use. We therefore sought to extend the platform’s applications to therapeutic GOF screens in the PF model.

To ensure high-fidelity mapping of genetic perturbations, we engineered constructs with a capturable barcode moiety in the 5’ untranslated region (UTR) followed by a coding sequence (*CMV-Barcode-Intervention-mGL-Sv40*) (**Fig 3A)**. This design minimizes discordance between the barcode and the functional intervention^33^. Similar to the prior *KO* screen, a pooled library of 10 putative fibrosis modulators, including ligands (*Wnt3a, Fgf9*, *Sst*, *Vash1*)^34–37^, enzymes (*Dio2, Mmp1b*)^38,39^, receptors/effectors (*Sstr2, Apcs*)^40,41^, and transcription factors (*Pparg, Klf15*)^42–44^ were delivered to the PF mouse model (**Fig 3B)**.

**Figure 3:**
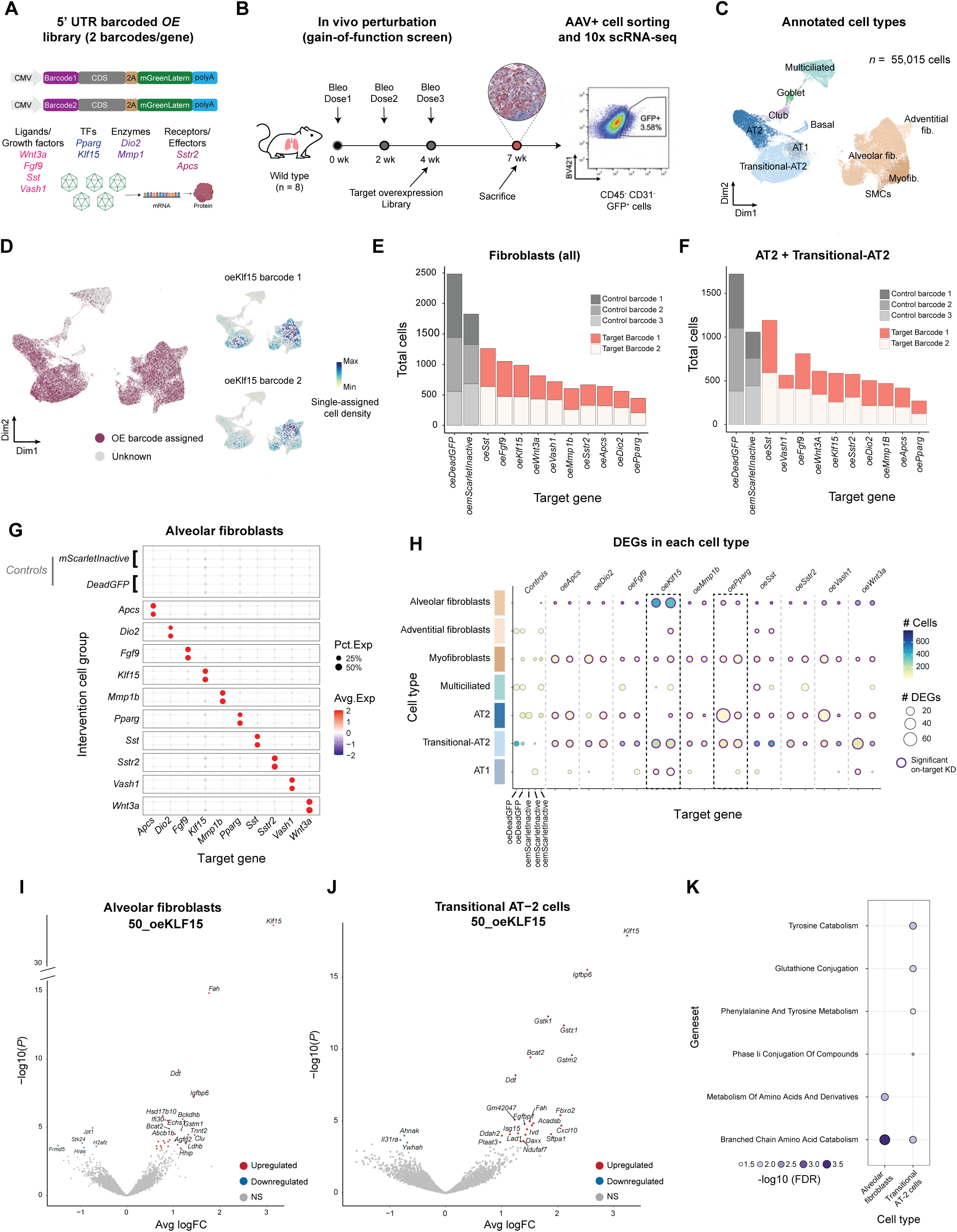
In vivo *GOF* mosaic screen reveals Klf15 as metabolic regulator of lung fibrosis resolution. **A.** Schematic depicting 5’ UTR barcoded overexpression (*OE*) construct design and target classes used for the GOF screen in PF model. **B.** schematic depicting the experimental *GOF* screen workflow, including PF model development, pooled library injection, FACS of transduced cells for scRNA-seq. **C.** UMAP of annotated cell types indicating mesenchymal and epithelial subcell-types. **D.** Distribution of assigned barcodes for *OE* (purple) and unknown (gray), with density plots showing cell-specific assignment of *Klf15* barcodes. **E.** Number of single barcode assigned fibroblast cells recovered per target genes and control. **F.** Number of single barcode assigned AT2 & transitional-AT2 recovered per target genes and control. **G.** Dot plot showing scaled expression of library genes in alveolar fibroblasts single-assigned to target and control barcodes. Dot size indicates the percentage of cells with detectable gene expression. Color scale indicates Z-score scaled average gene expression across assigned cell groups. **H.** Dot plot showing quantification of differentially expressed genes (DEGs) in each cell type in target genes assigned cells relative to deadGFP control cells (Adj *P < 0.1*). On-target (purple circle) indicates genes where the target gene is upregulated relative to *deadGFP* control cells. **I.** Volcano plots highlighting top differentially expressed genes by *Klf15* in alveolar fibroblasts (50 indicates barcode number), and **J**. transitional-AT2 cells. **K.** Gene set enrichment analysis for significantly upregulated genes in response to oe*Klf15 (FDR < 0.01*) in alveolar fibroblasts.

Following a three-week incubation period, we sorted GFP+ cells and recovered 55,015 single cell transcriptomes across the fibrotic lung epithelium and mesenchyme (**Fig3C-D Fig S3A-B**). Similar to the CRISPR KO screen, barcode capture was observed to be higher in alveolar fibroblasts and transitional-AT2 cells compared to other cell types (**Fig 3E-F, Fig S3C-D**). Notably, we observed that the number of single-assigned cells/target gene showed a significant negative correlation with the coding sequence (CDS) length, in both alveolar fibroblasts (Pearson r = −0.56*; P* = 0.003) and transitional-AT2 cells (Pearson r = −0.61; *P* < 0.001) (**Fig S3E-F**). This could be caused by lower rates of completed transcription for longer constructs and/or efficiency of AAV genome packaging^45^. Nevertheless, we recovered sufficient cell numbers for all targets to detect significant on-target expression (**Fig 3G, Fig S3G**). Downsampling analysis demonstrated high sensitivity for on-target *OE* detection dropping only under 100 cells (**Fig S3I**). This high sensitivity is attributable to the low basal expression of the target genes, which maximizes the signal-to-noise ratio of the ectopic transgene *OE* even with few cells.

Using a mutated, non-fluorescent GFP (deadGFP, EGFP Y67A) as an overexpression control, we performed differential expression analysis across cell types (**Fig 3H**). We identified the overexpression of Krüppel-like factor 15 (oe*Klf15*) as a potent metabolic switch in alveolar fibroblasts, driving branched-chain amino acid (BCAA) catabolism (*Bcat2, Bckdhb*), tyrosine degradation (*Fah*), and fatty acid fatty acid β-oxidation (*Echs1*) (**Fig 3I, K**). This shift toward high-efficiency oxidative metabolism was coupled with the repression of activation markers (*Hras, Jpt1*) and the upregulation of the anti-fibrotic Hedgehog inhibitor *Hhip*, known to preserve alveolar architecture and prevent myofibroblast differentiation by antagonizing Sonic Hedgehog (Shh) signaling^46–48^.

In transitional-AT2 cells, *Klf15 OE* promoted metabolic maturation through induction of amino acid metabolism and antioxidant defense (e.g., *Bcat2, Fah, Gstz1*) (**Fig 3J-K**) thereby redirecting the pathogenic cells towards a mature AT2 identity characterized by increased *Sftpa1* expression and the repression of the scaffolds *Ahnak* and *Ywhah* (**Fig 3J**). In contrast*, OE* of *Pparg* downregulated *Cdh1,* a gene essential for cell-cell adhesion in mature AT2 cells, and *Lad1* (gene involved in producing collagenous anchoring filament), and *Baiap2l1 (*gene involved in regulating actin cytoskeleton dynamics, cell shape, and motility) (**Fig S3H**). In contrast, Pparg effectively suppressed injury-associated genes such as *Fn1*, *Thbs1*, and *F3*, indicating its role as a repressor of pro-fibrotic signaling within the transitional-AT2 population^49,50^. Collectively, this data uncovers *Klf15* and *Pparg* as critical regulators in reestablishing metabolic homeostasis in both epithelial and mesenchymal compartments during fibrosis resolution.

### Enabling evaluation of perturbations’ therapeutic potential using disease- and human patient-relevant molecular features

While enrichment analysis successfully captured transcriptomic shifts of *Jak1*, *Tgfbr2*, and *Klf15*, other targets including *Pik3ca*, *Cebpb*, *Fgf9*, and *Wnt3a* produced more subtle transcriptional shifts in which effects could not be easily interpreted by looking at top DEGs, presumably due to the stochastic noise and cellular heterogeneity inherent in scRNA-seq data ^51,52^. To overcome these limitations and recover biologically meaningful effects from screens, we implemented a molecular feature-based framework that utilizes curated gene sets directly relevant to both the disease, tissue, and perturbation contexts (**Fig 4A**). Compared to GO or Reactome terms, these molecular features are, 1) derived from and/or validated against human patient sequencing datasets^53–56^, 2) pruned for gene-redundancy between features, and 3) mapped to encompass pathways that represent the range of known pathobiology and relevant physiological outcomes (e.g. fibrosis, regeneration, inflammation). By incorporating these elements, we aimed to move beyond open-ended gene lists to a functional ranking system that predicts the therapeutic efficacy and safety signals of diverse interventions *in vivo*.

**Figure 4.**
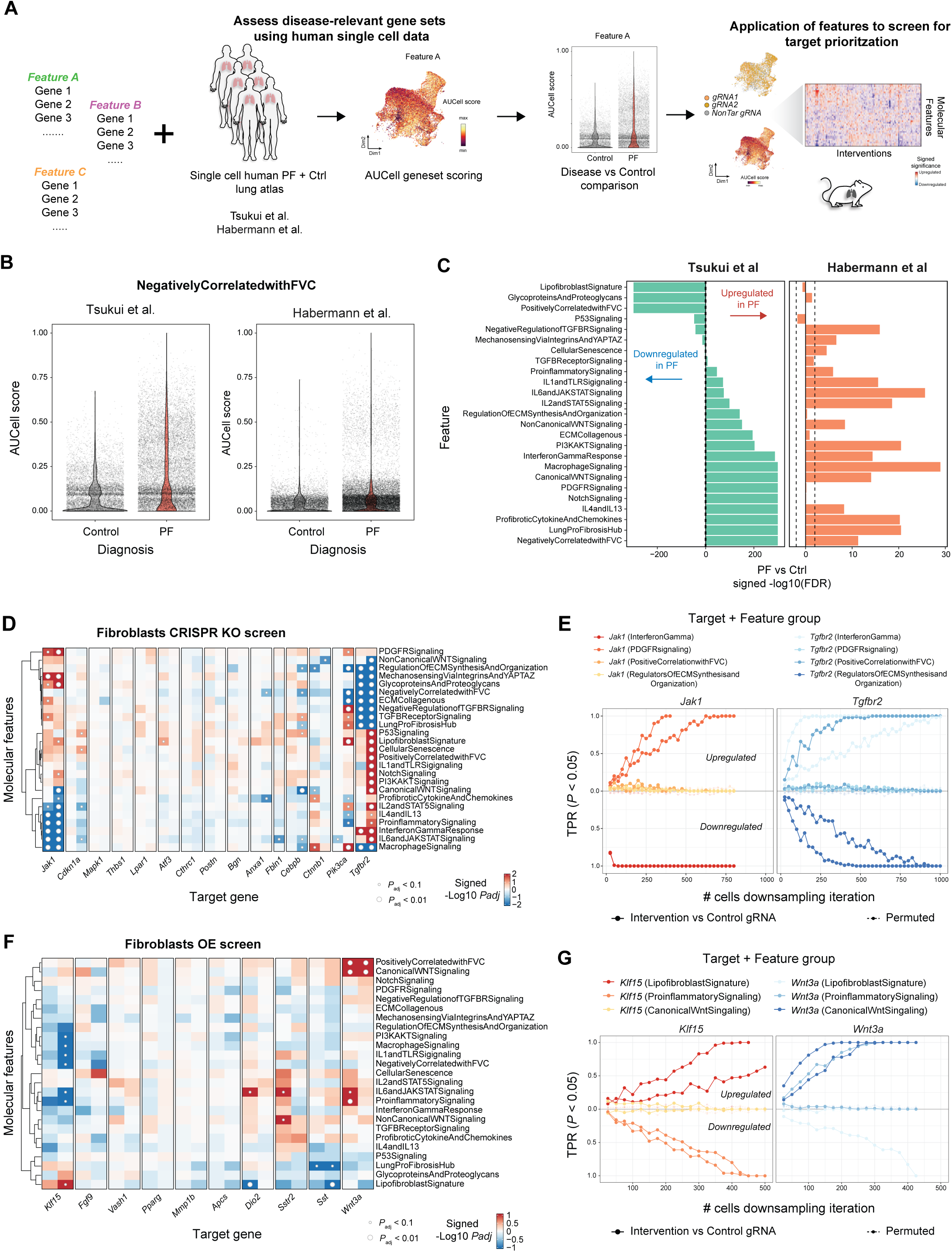
Transcriptomic analyses using disease-relevant molecular features enables prioritization across targets in PF screens. **A.** Schematic highlighting workflow for disease-relevant molecular feature curation and application for screen target assessment. **B.** Violin plots of single-cell AUCell scores for the *Negatively Correlated with FVC* feature shown for fibroblasts analyzed from two independent human scRNA-seq datasets Tsukui et al. (left) and Habermann et al. (right), grouped by study-designated diagnosis. **C.** Bar plot of PF vs healthy control effects (signed −log10 adj P value) for each disease feature (Wilcoxon test) for each dataset. **D.** Heatmap of target gRNA vs NonTar control molecular feature activity effects in CRISPR screen from Fig. 2. Color indicates signed −log10(*P*_adj_) from AUCell regression analysis of intervention vs control gRNA cells in fibroblasts. **E.** Sensitivity analysis for feature activity regression effects (related to Fig 4D) shown for different degrees of cell downsamping in *Jak1* and *Tgfbr2* KO cells relative to NonTar control cells. **F.** Heatmap of target *OE* vs deadGFP control molecular feature activity effects from screen in Fig. 3. Color indicates signed −log10(*P*_adj_) value from AUCell regression analysis of *OE* intervention vs control cell effects in fibroblasts. **G.** Sensitivity analysis for feature activity regression effects (related to Fig 4F) shown for different degrees of cell downsamping in *Klf15* and *Wnt3a OE* cells relative to deadGFP control cells. True positive rates (TPR; y-axis) in **E & G** were computed as the fraction of total sampling iterations (x-axis; *n*=100) that are significant (AUCell linear regression *P* < 0.05) in the same direction as observed at 100% (no downsampling) for a given feature (dotted lines represent effects from permuted label comparisons in each case).

The construction of these features involves ensuring specificity through a rigorous feature quality control (QC) pipeline (**Fig S4A**) that includes filtering gene sets by size, pathway relevance, overlap assessment, detection sensitivity based on overall gene expression (percent total UMIs) and co-expression levels when applied to mouse reference data (**Table S1**). The coordinated activity of genes in each QC-filtered feature can then be quantified (i.e. gene module scoring per cell, see Methods) and used to confirm disease relevance based on human scRNA-seq data, following which disease-specific features can be retained for perturbation effect assessment in screens (Fig 4A, see Methods). In addition to curated biological pathways, we integrated clinical features based on the directional correlation of gene expression with forced vital capacity (FVC). For example, *Positively Correlated with FVC* feature includes a set of genes that are downregulated as FVC declines in IPF. Conversely, *Negatively Correlated with FVC* feature include genes upregulated with FVC decline in PF^56–58^. We validated the utility of these features using independent IPF patient-derived scRNA-seq datasets^59,60^ (Tsukui et al.; Habermann et al.). AUCell scoring and differential testing between disease and control groups confirmed that disease-associated signatures such as the *Lung Pro-Fibrosis Hub* and *Negatively Correlated with FVC* were upregulated in IPF fibroblasts, while homeostatic signatures like the *Lipofibroblast Signature* were consistently suppressed (**Fig 4B-C**), thus confirming the disease specificity of these features.

Application of our molecular feature framework to the CRISPR *KO* screen successfully resolved both robust and subtle perturbations **(Fig 4D, S4B)**. *Tgfbr2 KO* induced a comprehensive suppression of profibrotic programs: *Tgfb Signaling, ECM Synthesis,* and the *Lung Pro-Fibrosis Hub,* while concurrently elevating broader inflammatory response including Interferon-γ (**Fig S4D),** consistent with its role in immune homeostasis. Conversely, *Jak1 KO* drove a distinct phenotypic shift toward a profibrotic fibroblast state, characterized by suppressed inflammatory features but increased *PDGFR Signaling*, mechanosensing (*Yap-Taz*), and non-collagenous ECM glycoproteins. Critically, while *Pik3ca* perturbation yielded negligible signal via standard DEG analysis, our framework unmasked a significant suppression of ECM organization coupled with induction of the *Lipofibroblast Signature* (*P_adj_* < 0.01) (**Fig 4D**). This shift indicates a successful conversion from a fibrotic to a homeostatic phenotype. Similar Wnt-regulatory roles were confirmed for *Cebpb* and *Ctnnb1* (**Fig 4D**). We observed similar, albeit non-significant, trends in transitional-AT2 cells (**Fig S4D**). To ensure these findings were not artifacts of technical noise or cell abundance, downsampling analysis confirmed the sensitivity of capturing significant effects for key features, with certain effects capturable with as few as 100-200 cells at over 80% sensitivity (regression *P* < 0.05, see Methods), validating the robustness of the observed biological outcomes for those gRNA-feature combinations (**Fig 4E**).

Next, our framework identified broad but directed shifts that e.g. DEG analysis or agnostic enrichment analysis might overlook (**Fig S4C**). *Klf15 OE* effectively reversed the fibrotic phenotype, significantly upregulating the *Lipofibroblast Signature* in fibroblasts and the *Positive Correlation with FVC* signature in transitional-AT2 cells, while simultaneously exerting an anti-inflammatory effect (**Fig 4F**). Notably, *Wnt3a OE* promoted the differentiation of transitional-AT2 cells toward a mature AT2 state, evidenced by increased *Surfactant Metabolism* and *Positively Correlated with FVC* signatures (**Fig 4F, Fig S4E**), effectively uncovering the effects that were failed to capture with standard pathway analysis. Interestingly, this Wnt3a driven repair was coupled with increase in pro-inflammatory features, suggesting a complex pro-repair and pro-inflammatory role for *Wnt3a* in established fibrosis^34,35,61^. As with the loss of function (*LOF*) effects, feature sensitivity for *Klf15* and *Wnt3a* reached *TPR >* 80% at modest cell counts (*n*∼300-400) confirming the scalability of the approach (**Fig 4G**). Collectively, these results demonstrate the capabilities of this orthogonal framework to establish a semi-quantitative ranking of interventions across a fixed set of pre-defined, human disease-relevant biological axes, which, can be further extended to other tissue and disease contexts following similar gene set curation and quality control workflows.

Collectively, these results demonstrate that our framework enables a semi-quantitative ranking of interventions across consistent biological axes. By utilizing non-overlapping gene sets, we can now compare disparate perturbations on a unified scale, a capability we next tested by extending the framework to different tissues, diseases, and species.

### Single-cell equine synovial landscape shares human-derived molecular features of osteoarthritis

Osteoarthritis (OA) represents another common class of age-related disease, characterized by cell death, inflammation, and catabolism^62^. To demonstrate the broad applicability and versatility of in vivo pooled screening, we conducted screens in horses with naturally occurring OA. Compared to murine models, the equine joint provides a highly relevant translational platform due to its comparable cartilage thickness, weight-bearing biomechanics, spontaneous disease onset, and chronic synovitis which closely mirror human OA^63,64,65^, thus making it an ideal clinical surrogate for therapeutic validation. We focused on the synovial tissue for our functional screens, given the critical role of chronic synovitis in OA-associated pain and progressive joint catabolism^66,67^.

To resolve the undercharacterized cellular landscape, we leveraged canonical human marker genes to annotate the equine synovium. We identified distinct fibroblasts populations including, lining *PRG4*^high^*CD55*^high 68^ and sublining fibroblasts (*PDGFRA*+*THY1*+) such as *CD34*^high^ and *CD34*^low^ subsets^69^ ^70^ (**Fig S5A–B**). Beyond these, the equine synovium revealed disease-associated fibroblasts including a unique lining subcluster, migratory fibroblasts characterized by high expression of *SOX5* and *AHNAK* (**Fig S5B-C**). These cell types and markers were independently validated in screening cohorts, confirming the robustness of the equine annotations.

Similar to PF, we curated molecular features of OA reflecting different pathophysiological states of the annotated synovial lining and sublining fibroblasts (**Table S2**). Diagnostic features of the diseased synovium included *Cartilage Matrix Degradation, TNFA Signaling,* and *IFNG Signaling*. Additionally, we curated precise mechanistic hubs to capture specific pathway architectures, including *JAK/STAT Activation, NFkB Activation* and the *NLRP3 Inflammasome*. Validation against human OA datasets^71^ (GSE216651) revealed conserved transcriptional signatures, including upregulation of inflammatory, fibrotic and metabolic stress features, confirming spontaneous equine OA as a high-fidelity translational surrogate (**Fig S5E-F**).

### miRNA knockdown screening identifies metabolic and inflammatory drivers in the equine synovium

We optimized viral gene delivery into OA synovium (see methods). While AAV-DJ and AAV-DJ9 were promiscuous, AAV5 demonstrated selective tropism for synovial lining and the *THY1*^high^*CD34*^low^ sublining subsets, along with transducing smooth muscle cells (**Fig. S5D**). Given its specificity, AAV5 was selected for intra-articular injection of vector libraries.

We then performed an in vivo *knockdown (KD)* screen using synthetic miRNAs targeting 21 genes implicated in OA (**Figure 5A**). Two weeks post-injection into OA horse fetlock joints, we recovered and sequenced 175,576 cells and assessed on-target knockdown in single-assigned synovial lining (*PRG4*^high^*CD55*^high^) and sublining fibroblasts (*THY1*^high^*CD34*^high^ and *THY1*^high^*CD34*^low^) (**Fig 5B**) which expressed *HTRA4* and *CXCL12*^72^, respectively, confirming pathological phenotype (**Fig 5C**). Statistical validation confirmed on target knockdown for nine targets (*LGALS3, COX7B, ROMO1, S100A11, COX6C, S100A6, S100A4, ADIRF, and CLEC3B)*, reaching significance in >40% of iterations (**Fig. S5G**). Our ability to detect on-target knockdown was significantly correlated with baseline expression levels of target genes (**Fig S5H**). Applying the molecular features, we found that *COX6C KD* significantly downregulated *Ion Channel* and *Oxidative Phosphorylation* features (**Fig 5E)**. Similarly, *S100A4* KD downregulated *Oxidative Phosphorylation,* while *LGALS3* KD yielded directionally consistent, albeit non-significant, trends on *Fibrotic ECM* and *Inflammatory Mediators,* demonstrating sensitivity to subtle in vivo effects. Together, these data establish a high-resolution functional KD screening pipeline in the equine joint, identifying *COX6C*, *S100A4*, and *LGALS3* as key metabolic and inflammatory regulators within the synovial landscape.

**Figure 5.**
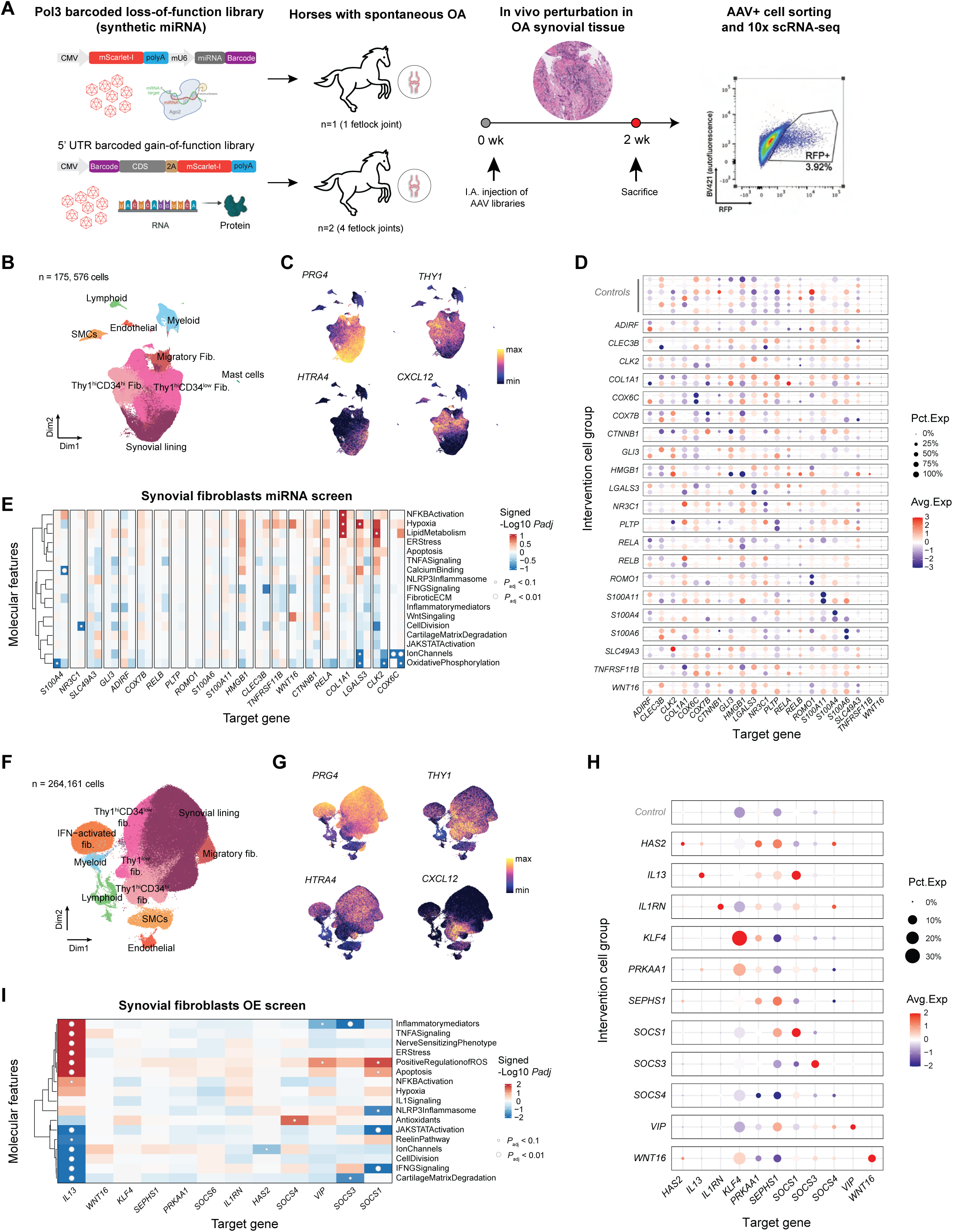
Equine *KD* and *GOF* screens identify molecular drivers of OA in the equine synovium. **A**. Schematic highlighting workflow for *KD* and *OE* screens in the synovial tissue of equine osteoarthritic fetlock joints. Fluorophore-positive cells were sorted after 2 weeks of expression and processed for scRNA-seq analysis. **B.** UMAP of synovial cells captured in miRNA screen from equine fetlock joints (*n*=1) colored by annotated cell type. **C.** UMAP visualization of cell type and disease-specific marker gene expression of synovial lining (*PRG4, HTRA4)* and sublining (*THY1, CXCL12*) fibroblasts in *KD* screen. **D.** Dot plot of scaled gene expression for library target genes among cell groups receiving each target miRNA or controls (scrambled miRNA) in lining and sublining fibroblasts. Dot size indicates the percentage of cells with detectable gene expression. Color scale indicates Z-score scaled average gene expression across assigned cell groups. **E.** Heatmap of target miRNA perturbation effects on molecular feature activity in pooled synovial lining and sublining fibroblasts. Color indicates signed −log10(*P*_adj_) from AUCell scoring of miRNA vs control cells per feature. **F.** UMAP of synovial cells harvested from equine fetlock joints (*n*=4) following *OE* screens colored by annotated celltype. **G.** UMAP visualization of cell type and disease-specific marker gene expression of synovial lining (*PRG4, HTRA4)* and sublining (*THY1, CXCL12*) fibroblasts in *GOF* screen. **H.** Dot plot of scaled gene expression for library target genes among cell groups receiving each target *OE* construct or control (mScarletInactive) construct across pooled synovial lining and sublining fibroblasts. **I.** Heatmap of target *OE* perturbation effects on molecular feature activity in pooled synovial lining and sublining fibroblasts. Color indicates signed −log10(*P*_adj_) from AUCell analysis of intervention vs control cells per feature.

### Equine GOF screening distinguishes the pleiotropic liabilities of IL13 from the targeted efficacy of SOCS factors

Complementing our knockdown studies, we performed in vivo *OE* screens of 12 candidate genes across immune (*IL13, IL1RN, SOCS1, SOCS3, SOCS4, SOCS6, VIP*), ECM (*HAS2, WNT16*), and metabolic (*PRKAA1, SEPHS1, KLF4*) modulators. From an initial pool of 264,161 captured cells, we focused on single-assigned synovial lining and sublining cells, which duly expressed *HTRA4* and *CXCL12* respectively (**Fig 5F-G**), with confirmed on-target *OE* (**Fig 5H, S5I**).

*IL13 OE* elicited the most profound and pleiotropic transcriptional shifts across molecular features (**Fig 5I**). Consistent with its known context-dependent roles, *IL13* significantly suppressed *JAK/STAT Activation*, *Ion Channels*, *IFNG Signaling*, and *Cartilage Matrix Degradation*, suggesting a protective, anabolic response. However, this was accompanied by a significant upregulation of *Inflammatory Mediators*, *TNFA Signaling*, *Nerve Sensitizing Phenotype*, *ER Stress*, *Positive Regulation of ROS*, *Apoptosis*, and *NFkappaB Activation* (**Fig 5I**), indicating concurrent activation of deleterious stress and inflammatory pathways.

In contrast, the Suppressor of Cytokine Signaling (SOCS) family members displayed more targeted anti-inflammatory profiles. *SOCS1*, a known negative regulator of JAK/STAT and interferon-gamma signaling (**Fig 5I**) ^73^ ^74^, significantly reduced *JAK/STAT Activation* and *IFNG Signaling* scores as expected. Furthermore, *SOCS1 OE* significantly suppressed the *NLRP3 Inflammasome* and *Inflammatory Mediators*, though it was associated with an increase in *Apoptosis* and *Positive Regulation of ROS*. Consistent with previous reports, *SOCS3 OE* produced anabolic and anti-inflammatory effects by significantly reducing *Inflammatory Mediators* and *Cartilage Matrix Degradation* signatures^75^, while *VIP* reduced inflammation and *SOCS4* uniquely increased the *Antioxidants* capacity (**Fig 5I**). Collectively, we validated the precise, pathway-specific efficacy of *SOCS* family members while uncovering the unappreciated pleiotropic liabilities of *IL13* thus functionally segregates safe, pathway-specific mechanisms from complex risks within a clinically relevant environment.

These results confirm that our pooled in vivo screens can interrogate effects on pathobiology and pathophysiology for both LOF and GOF perturbations in large animals, opening the door to multiplexed preclinical validation with superior scale and predictive validity for age-related diseases.

### Scaling mosaic screening for high-throughput discovery

Scalability of in vivo mosaic screening is inherently constrained by the cell numbers required for statistical power to discern phenotypic effects. Having established a sensitivity threshold of ∼250 cells per perturbation in prior screens, we sought to scale the platform to interrogate ∼300 constructs targeting 96 fibrosis-associated genes^53,76–80^ (**Fig 6A**). Similar to the earlier library design, we utilized a dual-gRNA architecture (3 constructs, 6 gRNAs per target gene plus controls) to ensure high-efficiency editing. Applying the same experimental workflow, we enriched transduced mesenchymal cells by negatively selecting the immune (CD45), endothelial (CD31), and epithelial (EpCAM) cells during FACS prior to single cell sequencing.

**Figure 6.**
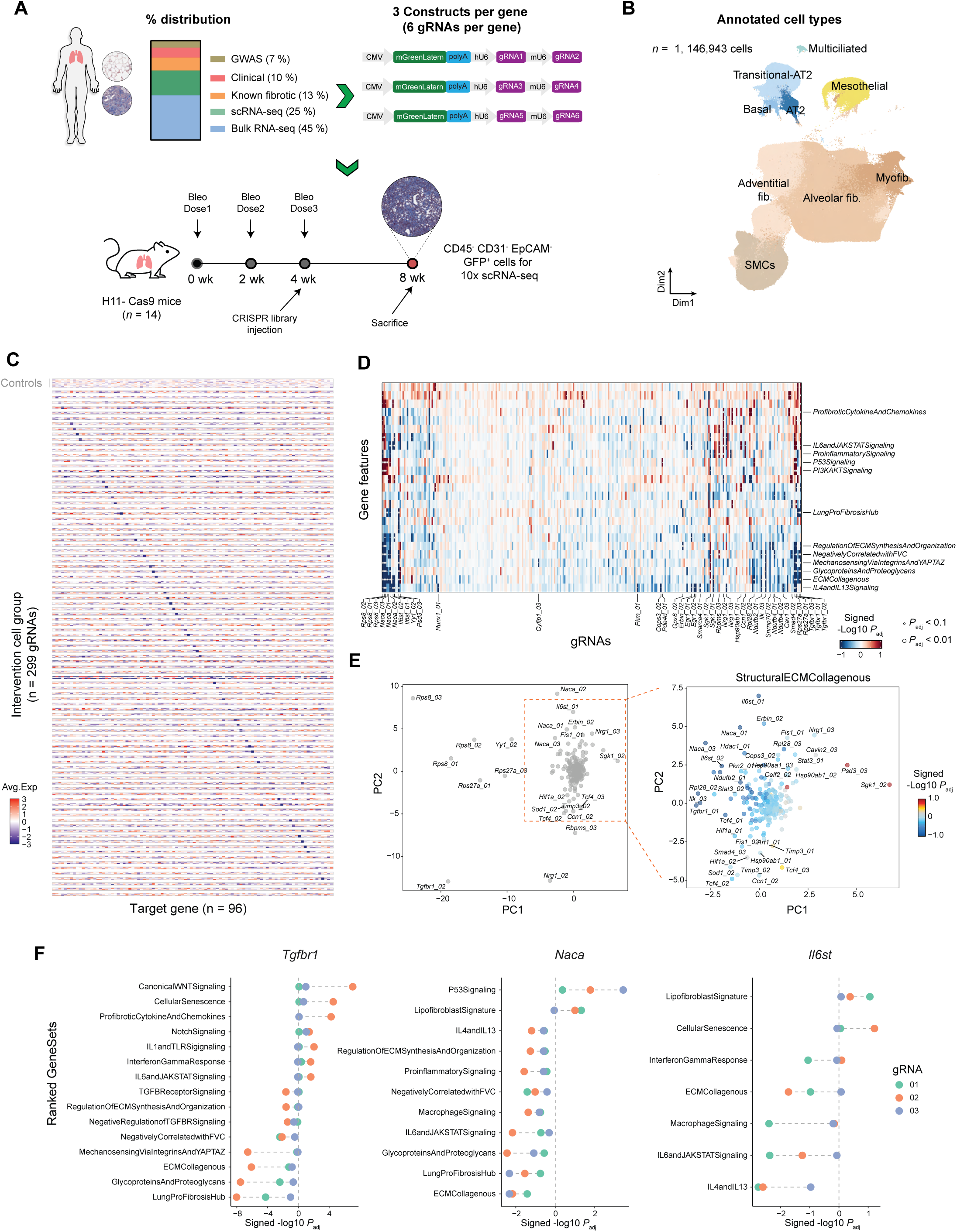
Large scale *in vivo* CRISPR *KO* mosaic screen identifies distinct anti-fibrotic and immunomodulatory targets. **A.** Schematic workflow for large-scale CRISPR *KO* mouse screen. Screen targets were included based on human genetics and transcriptomic, or clinical relevance from human PF patient data (*Top*), and three constructs with two gRNA each introduced into the experimental mosaic screen workflow. **B.** UMAP embedding of all cells (including single, multiple-assigned, and unassigned cells) colored by annotated cell type. **C.** Heatmap of scaled expression for library target genes across fibroblasts single-assigned to each gRNA construct. Color scale indicates Z-score scaled average gene expression across assigned cell groups. **D.** Heatmap of molecular feature activity effects across the large scale CRISPR screen. Color indicates signed −log10(*P*_adj_) from AUCell scoring of target gRNA vs NonTar control gRNA cells in single gRNA-assigned fibroblasts. Features where one or more gRNAs show effects at *P*_adj_ < 0.1 are highlighted, as are the corresponding gRNAs. **E.** PC biplot of the effects shown in D, highlighting gRNAs with the most effects across molecular features (left), and a zoomed in view of the same dimensionality reduction plot (right), with gRNAs colored by the *Structural ECM Collagenous* feature effect (signed −log10 *P*_ad_*_j_*). **F.** Waterfall plots of molecular feature effects (shown as signed −log10 *P*_adj_) for individual gRNAs targeting *Tgfbr1*, *Naca*, and *Il6st* are shown. Only features with gRNA vs inert control *P*_adj_ < 0.1 for at least one gRNA are shown, in each case.

We recovered over 1.1M cells across 14 mice, with >95% of cells identified as fibroblasts (**Fig 6B**). Comparison of gRNA-assigned cells against non-targeting controls confirmed consistent knockdown of target genes (**Fig 6C**). To analyze this extensive dataset, we used our previously described molecular features for a commensurable assessment of all targets’ anti-fibrotic and anti-inflammatory effects, identifying multiple constructs with significant effects (**Fig 6D**). Notably, we observed significant effects (regression *P*_adj_ < 0.01) on molecular features by at least one gRNA in 32 of the 96 targets tested (33%). The effects of perturbations were multi-faceted, with different targets affecting cellular phenotypes along multiple axes (**Fig 6E**), including some gRNA-specific effects related to extracellular matrix modulation and inflammatory signaling (Fig 6E). Notably, we are able to classify targets by specific effects, e.g. anti-fibrotic, even when targets had different mechanisms and overall transcriptomic effects. For example, gRNAs targeting *Tgfbr1* and the chaperone *Naca* both significantly reduce activity of *Lung Pro-Fibrosis Hub*, *ECM Collagenous*, and *Regulation of ECM Synthesis and Organization* features (**Fig 6F**), while showing opposite effects on inflammation and pathways associated with regeneration. gRNAs against *Il6st* had some anti-fibrotic effects while reducing activity of *IL6-* and *Macrophage Signaling* features (**Fig 6F**), consistent with the expected role as a universal signal transducing coreceptor for the Il6 family of cytokines. Thus this analytical approach provides a common ground for comparison and prediction even when the strongest effects of perturbing different target genes do not overlap. These results overall suggest 1) the reproducibility and scalability of our in vivo AAV screening technology; 2) the possibility to design and test libraries comprising several hundred gRNAs targeting a wide array of disease-relevant genes, and 3) an analytical framework that facilitates the identification of candidates eliciting therapeutically relevant transcriptomic effects, with the ability to further query nuanced effects and mechanisms of action.

### Orthogonal validation of screen findings for therapeutic efficacy

One of the most powerful uses for pooled in vivo screening would be to identify therapeutic targets able to reverse pathophysiology in human tissue. We therefore tested whether targets prioritized by our framework could reverse disease phenotypes in human primary ex vivo cultures.

Specifically, to verify hits that downregulated the *ECM Synthesis and Organization* feature in our two PF screens, we employed a human donor-derived precision cut lung slice (PCLS) ex vivo fibrosis model (n = 6-9 donor samples) (**Fig 7A**).

**Fig 7.**
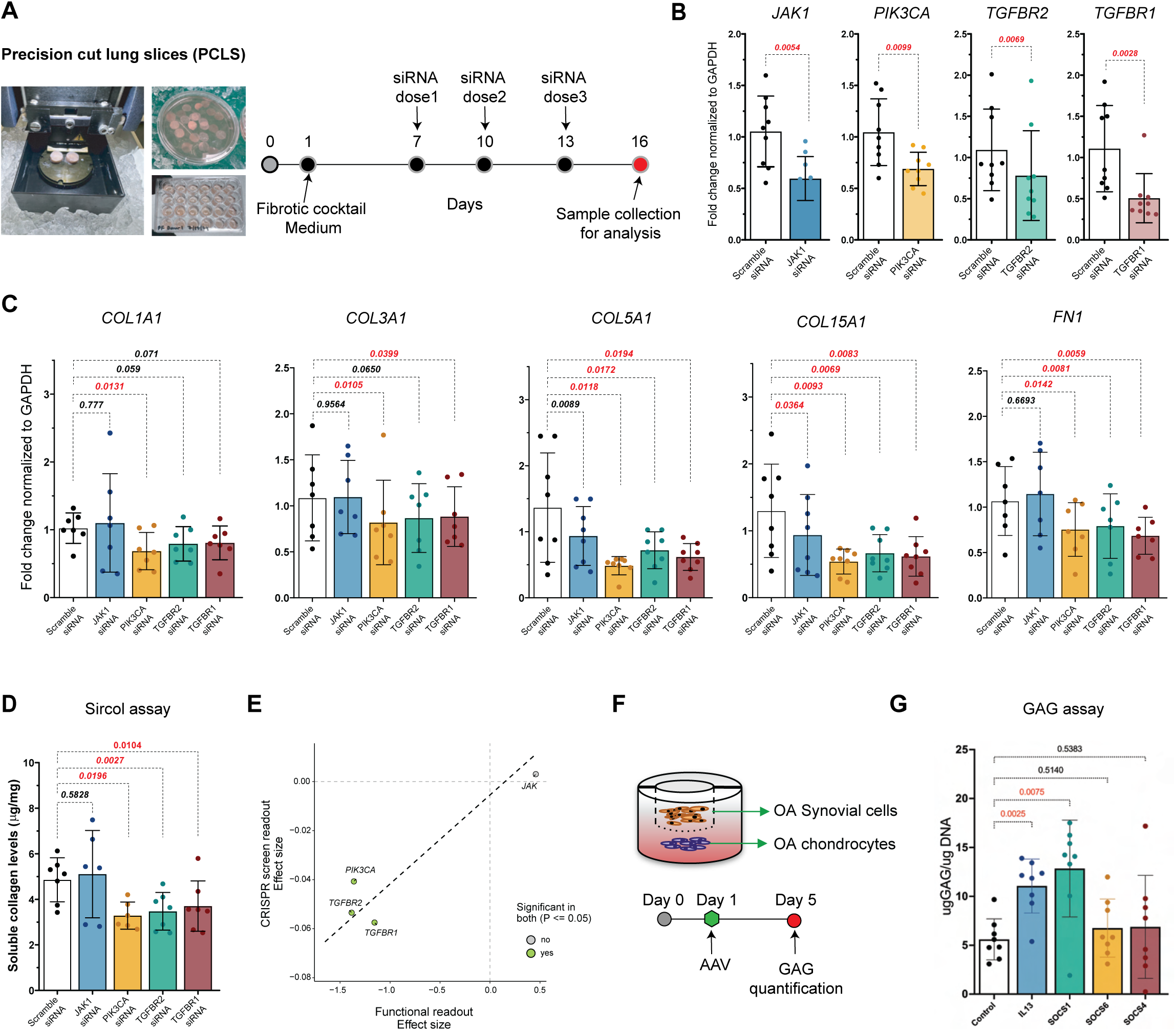
Validation of screen predictions in human ex vivo models. **A.** Schematic of human PCLS ex vivo model and siRNA knockdown workflow, including FC treatment and siRNA dosing. **B.** Bar plots showing target gene knockdown measured by RT-qPCR. **C**. Gene expression analysis by RT-qPCR. Fold expression change (normalized to GAPDH) of fibrotic markers (*COL1A1*, *COL3A1*, *COL5A1*, *COL15A1*, *FN1*) following siRNA knockdown (n = 7-9 donors per group). P value indicated on the top for respective comparisons. **D.** Quantification of soluble collagen levels (ug/mg) in sample treated with siRNAs targeting *TGFBR2* (*p* = 0.002), *TGFBR1* (*p = 0.010*), and *PIK3CA (p = 0.019)*, and *JAK1 (p = 0.58)* compared to control (n = 6-8 donors per group). Data sets in C & D are representative of mean ± standard deviation (Two-tailed paired T-test between scramble siRNA and target-specific siRNA groups). **E.** Scatter plot showing the correlation between observed effect size for ECM features in CRISPR screens and functional ECM readout by sircol assay. **F.** Schematic of human OA patient derived co-culture model and saturating AAV *OE*. **G.** Quantification of GAG content (µg GAG/µg DNA) in human OA chondrocytes co-cultured with patient-matched OA synovial cells treated with AAV expressing IL13 or SOCS1 or SOCS6 or SOCS4 vs untreated control at Day 5 (n = 8 donors). Data represented here as mean ± standard deviation (One-way Brown-Forsythe ANOVA with Bonferroni correction, P<0.05).

One of the most powerful uses for pooled in vivo screening would be to identify therapeutic targets able to reverse pathophysiology in human tissue. We therefore tested whether targets prioritized by our framework could reverse disease phenotypes in human primary ex vivo cultures. Specifically, to verify hits that downregulated the *ECM Synthesis and Organization* feature in our two PF screens, we employed a human donor-derived precision cut lung slice (PCLS) ex vivo fibrosis model (n = 6-9 donor samples) (**Fig 7A**). Treatment of donor tissue-derived PCLS with fibrotic cocktail medium successfully recapitulated the fibrotic phenotype, as confirmed by soluble collagen production and Gene Set Variation Analysis (GSVA) scoring of *Lung Pro-Fibrotic Hub* and *ECM Synthesis and Organization* signatures (**Fig S7A-C**). Targeting candidate genes via siRNA revealed distinct functional outcomes. Knockdown of *PIK3CA*, *TGFBR2*, and *TGFBR1* significantly reduced key fibrotic markers (*COL1A1*, *COL3A1*, *COL5A1*, *COL15A1*, and *FN1*) and soluble collagen production (*p < 0.02*) (**Fig 7B-D**). In contrast, *JAK1* knockdown did not yield significant reductions (*p = 0.58*), distinguishing structural drivers from broad immunomodulators. A scatter plot suggests a linear correlation between functional effect size in human PCLS and the initial CRISPR screen readout (**Fig 7E**).

Next, to validate equine screen hits identified through *OE* screen, we utilized a patient-derived co-culture model (n=8 biological donors) to recapitulate the pathological crosstalk where inflamed synovium drives glycosaminoglycan (GAG) loss in chondrocytes (**Fig 7F**). We selected targets representing a range of phenotypic potencies, including *IL13* and *SOCS1* due to their profound effects on inflammatory and cartilage-degrading features, and *SOCS4* and *SOCS6* as comparators with weaker effects **(Fig 5G)**. Consistent with screen observations, viral delivery of *IL13* and *SOCS*1 at a saturating dose significantly restored chondrogenic function as a measure of total sulfated GAG content (*p = 0.0025* and *p = 0.0075*) relative to controls (**Fig 7G**). This anabolic recovery confirms distinct screen-defined profiles: IL13 mitigated matrix degradation, while SOCS1 dampened broader inflammatory drivers, including the NLRP3 and JAK/STAT pathways (Fig 5I)^81,82^. In contrast, *OE* of *SOCS6* (*p=0.5140*) and *SOCS4* (*p=0.5383*) failed to significantly alter GAG content (**Fig 7G**), validating their classification as weaker modulators in the primary screen. Collectively, the concordance observed between in vivo screen signatures and human ex vivo functional outcomes demonstrates the robustness of our target validation pipeline across animal models and human biology.

## Discussion

Extending high-throughput functional genomics to intact diseased tissues requires solving a set of challenges. The first is delivering varied payloads to organs with pathophysiological changes^83^. To overcome the inefficiency of transduction in fibrotic and degenerated tissues, we performed a systematic exploration of AAV receptor tropism. Using a barcoded AAV library, we empirically identified vectors with optimal performance in fibrotic lung epithelium and fibroblasts, and osteoarthritic equine joints, enabling efficient targeting of pathologically relevant cells and ensuring sufficient transduced cell numbers to support the required screening library complexity. Second, we demonstrated perturbations spanning targets relevant across drug modalities, including *GOF* and *KD*, without relying on difficult-to-deliver CRISPR tools. This is particularly critical for chronic diseases where pathology may be driven by haploinsufficiency or the loss of homeostatic gene expression, necessitating *GOF* strategies for therapeutic correction. Beyond balancing gene expression, this also allows for the screening of novel protein variants and therapeutic peptides.

Another major challenge of high-throughput screening is discerning therapeutically relevant signals, especially in heterogeneous disease tissues with diverse cell states^51,84^. We addressed this by complementing differential gene expression and common pathway enrichment analysis options with a framework of patient-derived molecular features linking transcriptional changes directly to disease-specific pathophysiological functions such as ECM organization, lung function, and homeostatic fibroblast states. This framework provides a more comprehensive readout on how a perturbation impacts disease pathology. For example, this approach enabled us to visualize how interventions like *Tgfbr2*, *Pik3ca*, and *Klf15* downregulate profibrotic signatures while upregulating homeostatic markers. Crucially, our power analysis revealed that these curated features yield high sensitivity with saturation at ∼200–250 cells per target, confirming that in vivo screens can scale to large gene sets with sufficient statistical power, as demonstrated in our lung fibroblast screen (Fig 6).

A defining advance of this work is the extension of in vivo screening to large animal models. This opens up more translationally relevant models, such as naturally occurring disease with human-like etiology, and subjects with varied genetics and life history. It also enables the investigation of biological processes poorly recapitulated by standard murine models, for genetic and/or anatomical reasons. For example, rodent models of OA often fail to recapitulate human OA pathophysiology, due to profound differences in joint size, cartilage thickness, and biomechanical loading. By applying the platform in the clinically relevant niche of spontaneously occurring OA in horse joints, we identified *IL13* and *SOCS1* as potent regulators of inflammation and cartilage degradation. To our knowledge, this study represents the first application of in vivo transcriptomic screening in any large animal model, establishing a foundation for high-throughput single cell functional genomics in non-rodent species. We closed the translational loop by validating predictions from molecular features in human ex vivo models (PCLS and knee joint tissue). Strong correlation between our mosaic screen observations and human tissue physiological responses confirms that this platform can predict potentially therapeutic outcomes through multiple biological mechanisms.

Despite its utility, in vivo mosaic screening in its current state has several limitations: First, because mosaic perturbation intentionally leaves most of the tissue unperturbed, organ size and cellularity puts limits on the number of perturbations that can be multiplexed in a single animal. Since discerning transcriptome signals requires only a few hundred cells per target (Fig 4), this is most relevant for small glands, sensory organs, and sub-regions of e.g. the brain. This can be mitigated by using large animal models, as illustrated by our ability to screen 48 constructs in the synovium of a single equine joint. Second, while our data shows that significant effects can be detected for both cell-autonomous and paracrine signaling targets, the latter are likely mediated by interactions in the local niche. Predicting effects involving organ-to-organ endocrine signaling would likely require paired datasets linking endocrine signaling phenotypes to transcriptomic cell states. Finally, the three perturbation modalities utilized here primarily modulate transcriptional regulation, which may yield false negative results for targets predominantly regulated posttranslationally. Though not demonstrated here, the platform could be adapted to perturb at the protein level, by delivering different coding sequences such as proximity inducers or nanobodies.

In conclusion, we described a mosaic screening platform for modality-agnostic validation of therapeutic targets at scale, by assessing causal in vivo perturbation effects using disease-relevant molecular features based on patient-derived datasets. By enabling delivery and assessment of multiple perturbation modalities across pathological tissues in rodent and large animal models, this platform provides a powerful strategy for dissecting the multicellular molecular circuits driving chronic diseases and accelerating identification of novel therapeutic targets.

## Author contributions

V.S. conceived, designed, and supervised in vivo CRISPR mosaic screens; co-designed and supervised gain-of-function screens in mice; curated PF molecular features; optimized protocol for lung single cell isolation; performed human PCLS studies and analyzed data. V.K. performed all computational analysis of screen data. N.S. conceived, co-designed gain-of-function and miRNA-based equine screens, curated molecular features and analyzed data. D.R.F. designed barcoding strategy for gain-of-function screens and performed molecular biology work. L.C., C.T. designed, produced, and performed serotype screens. H.M., J.C., S.S., A.L. performed tissue dissociations, FACS, and scRNA-seq. I.D. built the cellranger and perturbation assignment pipelines. D.P., K.D. demultiplexed sequencing data and processed perturbation assignment pipeline. A.G, A.V., G.D. performed data processing and analysis. L.F. designed barcoded libraries for miRNA screens and performed molecular biology work. S.H., J.C.S performed vector library preparation. K.F., C.B, D.G., produced and purified AAV vectors and performed quality control. J.N. performed in vivo lung experiments, processed tissue samples and assisted with PCLS. C.L. performed RT-qPCR and sircol assays. A.B. B.B.P., and R.W.L. supervised and performed equine studies. F.L. provided support with resources. M.B.J. conceptualized and supervised the work. V.S., V.K., N.S., and M.B.J. co-wrote the manuscript, with input and review from all authors.

## Acknowledgements

We thank Dr. Toby Maher for advice and fruitful discussions on work related to pulmonary fibrosis. We thank Dr. David Hunter for advice and fruitful discussions on work related to osteoarthritis. We thank Dominic Grun, Randy Platt, Wyatt Morgan, Atray Dixit for advice on pooled CRISPR screening. We thank all of Gordian’s investors for their support. We thank the University of Florida for sourcing equine subjects for this study. We thank the tissue procurement agencies for sourcing human tissue samples. NIH SBIR 6R44ES032515-02 helped fund human data for molecular features.

## Methods

### Animals

Mouse strains, wildtype (strain# 000664) or H11^Cas9^ CRISPR/Cas9 knock-in (B6J.129(Cg)-*Igs2^tm1.1(CAG-cas9*)Mmw^*/J, strain# 028239) used in this study were sourced from Jackson laboratory. All mouse experiments were approved by Gordian Institutional Animal Care and Use Committee (IACUC). All animals were handled in accordance with the IACUC committee guidelines for humane care and use of laboratory animals. Mice cages, bedding, food and water were autoclaved and supplied with fluids as needed during the experimental period. Animals were maintained on the 12h light/dark cycle and monitored daily for health. Lame thoroughbred geldings (3–5 years old, 450–500 kg) with radiographic evidence of osteoarthritis were enrolled in this study under IACUC approval from the University of Florida. All horses exhibiting mild-to-moderate osteoarthritis (OA) in both the right and left fetlock joints were enrolled in the study for OA studies.

### Human samples

Excised sub-transplant quality lung biospecimens from post-mortem donors were obtained from donor procurement agencies including, National Disease Research Interchange (NDRI), Donor Network West (DNW), or The International Institute for the Advancement of Medicine (IIAM) under research authorization agreement. Surgical discards of cartilage and synovial tissues were procured from patients undergoing total knee arthroscopies via NDRI.

### AAV serotype library production

Individual single stranded (ss) AAV serotypes were packaged with barcoded DNA pools under pol3 promoter (ITR-CMV-mGreenLantern-Sv40(polyA)-Pol3-Barcode-ITR) separately. A total of 13 AAV serotypes (AAV1, AAV2, AAV2.5, AAV3, AAV5, AAV8, AAV9, AAV-DJ, AAV-DJ8, AAV-DJ9, AAV-rh10, AAV-NP40, AAV-NP59) were produced for screening in horses and 16 serotypes (AAV1, AAV2, AAV2.5, AAV3, AAV5, AAV8, AAV9, AAV2.5, AAV-8P2, AAV-DJ, AAV-DJ8, AAV-DJ9, AAV-rh1, AAV-rh10, AAV-NP40, AAV-NP59) were produced for screening in mice. AAV vectors were produced using transient transfection in HEK293T cells as previously described^85^. Briefly, HEK293T cells were transfected using a 1:1 PEI to DNA ratio with a DNA pool consisting of a mass ratio of 2:1:1 pHelper:rep-cap:GOI plasmids. 3 days post-transfection, the cells and media were harvested, treated with benzonase to remove excess DNA, PEG-precipitated, and processed for purification by iodixanol ultracentrifugation.

Each AAV was quantified by vector genome titering using hydrolysis probe-based qPCR where primers and probes bind and amplify the conserved intronic or barcode promoter region of the vector genome. Vector genomes were extracted from capsids using the Qiagen Minelute Virus Spin Kit and the genomes were qPCR-amplified using the cycle conditions per manufacturer instructions for the Applied Biosystems Taqman™ Gene Expression Master Mix (Cat# 4369016 Thermo Scientific): 50°C for 2 min, 95°C for 10 min, followed by 40 cycles of 95°C for 15 s and 60°C for 1 minute with a fluorescence scan. For each serotype pool, an equimolar amount of AAV was pooled prior to injection.

### AAV serotype screening in mice and horses

For serotype screening targeting epithelial and fibroblasts in mice, a pool of 16 serotypes were intratracheally injected in bleomycin induced fibrosis (n=5) and allowed to express for 4 weeks. After 4 weeks, tissues were harvested for analysis to determine the barcode enrichment in fibroblasts and epithelial cells following mice tissue dissociation and single cell protocol.

For serotype screening in chondrocytes and synovial fibroblasts, a pooled library of 13 AAV serotypes encoding a GFP reporter was delivered intra-articularly into the right and left fetlock joints of horses (n=2) and allowed to express for 4 weeks. At the endpoint, chondrocytes and synovial cells were harvested and GFP-positive cells were enriched by flow cytometry. Enriched populations were then analyzed to quantify serotype-specific barcode representation in fibroblasts and epithelial cells following tissue dissociation and single-cell processing.

### gRNA library design and AAV production

We designed four to six gRNAs per target gene against the Mouse GRCm38 genome (Ensembl v.102) using the Broad Institute’s Crispick CRISPRko selection tool. To minimize steric hindrance, gRNA binding sites were spaced at least 40 bp apart. For vector construction, gRNAs were synthesized by GenScript and cloned into an AAV plasmid backbone. Each vector contained two gRNAs driven by a human U6 and either a bovine or mouse U6 promoter, alongside a CMV promoter driving mGreenLantern or mScarlet expression with an SV40 polyadenylation signal. Perturbations were identified via direct capture of the gRNA sequences, which served as unique barcodes. All gRNA sequences used in this study are listed in **supplemental file 1**.

Individual DNA constructs were pooled at an equimolar ratio before AAV production. AAV-DJ9 vector libraries were produced using HEK293T cells as described previously. Briefly, 1:1 PEI to DNA ratio with a gRNA library pool consisting of a mass ratio of 2:1:1 pHelper:rep-cap:GOI plasmids. 3 days post-transfection, the cells and media were harvested, treated with benzonase to remove excess DNA, PEG-precipitated, and processed for purification by iodixanol ultracentrifugation. Each AAV was quantified by vector genome titering using hydrolysis probe-based qPCR method.

### Overexpression and miRNA libraries

For mouse and horse *OE* constructs, the barcoded coding DNA sequence (CDS) for each target gene was synthesized by Genscript and cloned into a CMV promoter plasmid backbone with either monomeric(m)GreenLantern or mScarlet-I fluorophore. For horse miRNA constructs, miRNA hairpins were designed against target gene mRNA transcripts using the Broad Institute’s RNAi Consortium (TRC) shRNA design rule set algorithm. Barcoded miRNAs were synthesized by Genscript and cloned into a dual CMV and murine U6 promoter plasmid backbone with either monomeric(m)GreenLantern or mScarlet-I fluorophore. To prepare endotoxin free DNA for AAV production, individual mouse and horse *OE* plasmid constructs were transformed into NEB® 5-alpha Competent E. coli (Cat# C2987H, New England Biolabs), cultured in LB broth (Cat# L8000, Teknova), and purified with either the I-Blue Mini Plasmid Endo-Free Kit (Cat# IB47176, IBI Scientific) or EndoFree Plasmid Maxi Kit (Cat# 12362, Qiagen). For the horse miRNA library, individual horse miRNA plasmid constructs were prepared directly by Genscript as research grade endotoxin free (≤0.1 EU/μg endotoxin) DNA preps. For the final library injection mix, individual construct DNA preps were pooled equimolar for miRNA libraries or according to a specified ratio adjusting for AAV packaging efficiency for *OE* libraries. For library representation analysis, the final DNA injection mix is sequenced by a nanopore-based DNA plasmid sequencing service, Whole Plasmid Sequencing (Quintara Bio) or Plasmid-EZ (GENEWIZ), and the generated FASTQs were used for analysis. Post plasmid QC, AAV vector libraries were produced using HEK293T cells as described in the previous section.

### In vivo mosaic screening in mice model pulmonary fibrosis

Persistent pulmonary fibrosis was induced in wildtype C57BL/6J or Cas9 (B6J.129(Cg)-*Igs2^tm1.1(CAG-cas9*)Mmw^*/J) mice by intratracheal injection of bleomycin (1 mg/kg) bi-weekly for a total of 3 doses as described previously^9^. 2e11 vg of AAV-DJ9 gRNA or *OE* library was mixed with 100 uM doxorubicin (Cat# D1515, Sigma) to enhance transduction, and injected along with the final dose of bleomycin. Perturbations were allowed to express for 3 (GOF screen) or 4 weeks (CRISPR screens) before lungs were harvested for analysis.

### Mice lung single cell isolation and FACS

Mouse lung tissue dissociation and FACS were performed as described previously^86^. Briefly, the body was perfused via right ventricular cannulation to preferentially clear blood from the lungs, with downstream systemic clearance. Lungs were then inflated with digestion buffer containing Collagenase type I (450 U/ml (Cat# 17100017, Thermofisher), Dispase II (5 U/ml) (Cat# 04942078001, Sigma) and DNase I (0.33U/ml) (Cat#10104159001, Sigma) in DMEM/F12 (Cat# SH30023.01, Cytiva). Lung tissue was excised and minced with blade and incubated at 37 degrees for 30 min. Enzymatic activity of the buffer was neutralized using 10% FBS containing DMEM/F12 medium. Tissue homogenate was then filtered through a 100 µm strainer, followed by RBC lysis. Cells were pelleted by centrifugation at 300 × g for 2 minutes at 4°C. Cells were resuspended in 0.04% BSA buffer with antibodies against CD45 (Cat# 130-052-301, Miltenyibiotech), CD31 (Cat# 130-097-418, Miltenyibiotech) and EpCAM (Cat# 130-105-958, Miltenyibiotech), Biolegend to deplete immune and endothelial cells using MACS protocol (Cat#130-042-401, Miltenyibiotech) as per manufacturers instructions. CD45, CD31, and EpCAM depleted live cells were FACS sorted for transduced GFP positive cells at a final concentration of 1,000 cells/µl.

### Screening in Horse spontaneous OA model

Pooled libraries were generated using AAV-NP59, selected for its selective transduction of the equine cartilage, and AAV5 for its synovial tissue tropism, using a suspension-based production system. Lame thoroughbred geldings were screened for serum neutralizing antibodies against AAV-NP59 and AAV5, and none were detected. In one horse (5 year old gelding), the right fetlock joint was injected intra-articularly (1 mL per joint) with 5 × 10¹¹ vg of a barcoded AAV5 library encoding a pooled set of microRNAs targeting 21 individual genes, driven by a CMV promoter and coupled to an mScarlet-I fluorophore. In the two additional 3 year old horses (four fetlock joints), each joint was injected with 5 × 10¹¹ vg of a barcoded AAV5 library expressing pooled *OE* cassettes for a different set of 18 genes, also under a CMV promoter with an mScarlet-I reporter. Transgene expression was allowed to proceed in vivo for two weeks, after which synovial tissues were harvested and immediately placed in HypoThermosol supplemented with 45 μM actinomycin D (Cat# HB1477, Hello Bio) until single-cell processing.

### Horse cartilage and synovium single cell isolation and 10x sequencing

Cartilage and synovial tissues were finely minced prior to enzymatic dissociation. Cartilage fragments were first incubated in pronase (10 mg/mL, DMEM/F-12 supplemented with 6 μM actinomycin D) for 30 min at 37 °C, centrifuged at 300 × g for 5 min, and subsequently digested in collagenase II (4 mg/mL, DMEM/F-12 supplemented with 6 μM actinomycin D) for 30 min at 37 °C. Synovial tissue fragments were digested in collagenase II (4 mg/mL) for 1 h at 37 °C. Following digestion, both cartilage and synovium derived cell suspensions were treated with 1× RBC lysis buffer for 4 min at room temperature, washed, and counted prior to downstream applications. Live RFP positive cells were FACS sorted for the subsequent 10x loading.

### Sequencing of intervention and single cell gene expression libraries

Single-cell capture was performed with 10x Genomics’ Chromium Single Cell 5’ Kit v2 or v3 with a spike-in of 50 pmols of relevant capture-sequence oligo(s) to the RT master mix. Gene expression libraries were prepared following the manufacturer’s protocol. To recover intervention tag molecules, the supernatant from the cDNA amplified SPRI selection was further purified with additional 1.2x SPRI selection. 20ng of the resulting DNA was used as a template, and amplified using Q5 High-Fidelity DNA Polymerase according to the following protocol: (1) 98C for 30 s, (2) 20 cycles of 98C for 10 s, 65C for 30 s, 72C for 60 s, (3) 72C for 10mins. The resulting intervention-tag library was purified using double-sided 0.6X-1.2X SPRI selection. Both the gene expression libraries and their corresponding intervention-tag libraries were sequenced using the NextSeq 550 75-cycle kit or NovaSeq X 300-cycles kit with the following cycle distribution: 28 to Read1, 8 to Index, and 56 to Read2 for NextSeq, and 10 each on Index1 & 2, and 150 each on Read1 & 2 for NovaSeq.

### scRNA-seq data processing and analysis

#### 10x single cell RNAseq read alignment and adaptor removal

All samples were run using fastp version 1.0.1 (https://github.com/OpenGene/fastp) with quality settings --qualified_quality_phred = 20 and --unqualified_percent_limit = 40. For sequences with R2 over 90 reads were trimmed to a max of 90bp. For reads with R2 of 56bp, sequences were not trimmed, just quality filtered. All samples were run using Cellranger version 9.0.1 against a pre-built mouse genome reference refdata-gex-mm10-2020-A.tar.gz from 10X Genomics and an Ensembl reference 106 for Equus caballus (Horse), the latter built using the mkref command from cellranger.

### Intervention barcode read alignment and quantification

Perturbation reads were assigned by splitting R1 into cellranger cell barcode and UMI based on a 10x sequencing version (16bp cell barcode and 12bp UMI for current versions). A dataframe with 4 columns (10x cell barcode, 10X UMI, R2, and read count) is used to assign each unique R2 to a perturbation using the unique barcode for each perturbation. The first 10,000 reads of each R2 fastq file were sampled to determine the fixed offset values for the barcode. These fixed positions were then used to do a 1 Hamming error match and all unique R2 sequences. Then the cell barcodes were corrected to the 10X cell barcode whitelist with a Hamming distance of 1. Only valid 10x cell barcodes were kept. Then the R2 sequences were dropped and the reads for each unique cell barcode, UMI, perturbation were collapsed. A read filtering step was performed by determining the threshold for background read counts based on the distribution of reads per UMI. Low count (<∼10) reads are then discarded. After read filtering, a matrix of UMI counts per cell barcode per perturbation is calculated.

### Intervention barcode assignment to single cells

In order to assign interventions to single cells, we implemented a probabilistic framework adapted from the beta-binomial model approach used by scDemultiplex algorithm for demultiplexing HashTag Oligo (HTO) counts^87^. Similar to HTO data, AAV library pooled barcode UMI counts exhibit overdispersions that do not always follow a poisson or a simpler binomial process. For each screen type (*KO*, *OE,* or *KD*), we directly use empty 10x droplets (defined as droplets having total cDNA UMIs < 300) to model our ambient fraction per 10x lane. After generating the cell x intervention barcode UMI count matrices (see section Intervention barcode read alignment and quantification above), we fit a beta-binomial (alpha, beta) model given the observed barcode-specific UMI counts in the cell for each intervention relative to empty-droplets from the same 10x lane. This directly estimates expected background proportion and overdispersion from experimentally observed contamination that can be a consequence of ambient RNA or overamplification. For each non-empty droplet (QC-filtered cell) and guide, we thus compute an upper tail beta binomial *p*-value conditioned on the total guide UMI capture for that droplet given the expected background. The assignment p-values are then adjusted per perturbation across filtered cells in the 10x lane using Benjamini–Hochberg correction. Interventions were independently assigned to droplets with adjusted assignment *P* < 0.01. For *OE* and miRNA screens, if this resulted in more than 1 barcode assigned to the same cell, this constituted a Multiple assigned cell. For the CRISPR screens, which utilized dual-gRNA constructs, guide assignments from the human U6 and mouse or bovine U6 promoters were evaluated independently. Cells were classified as single-assigned when both promoter-derived assignments corresponded to the same AAV construct. If an assignment was obtained from only one promoter, the cell was assigned to that AAV construct. In cases where promoter-specific assignments were discordant (i.e., corresponding to different AAV constructs), the cell was classified as “Multiple” assigned. In all cases, if no assignment was detected at Adj *P* < 0.01 they were called as “Unknown”. Only single-assigned cells were retained for subsequent (*KO*, *OE* or *KD*) perturbation effect analyses.

### scRNA-seq quality control, dimensionality reduction, clustering and cell type annotation

Cells were filtered for a minimum and maximum total UMI and number of genes detected across all samples specific to a given screen, maximum percent total mitochondrial reads (5%) and filtered for 10x doublets using DoubletDetection package in Python (http://doi.org/10.5281/zenodo.2678041). Filtered cells raw gene expression counts were then run through SCTransform normalization (SCT-normalized) using the Seurat R package, following which PCA and UMAP were used to embed cells in lower dimensional space. Harmony was used to adjust for animal effects where present ^88^. For lung screen scRNA-seq data, filtered cells were annotated using a combination of reference-based annotation using ACTINN ^89^ against the human cell lung atlas^90^. Cell type labels were assigned to louvain clusters based on consensus (majority vote) cell type labels from reference-based ACTINN prediction, together with inspecting cell type-specific markers. For horse screens, previously identified synovial cell marker genes from human synovial tissue single cell sequencing were used to assign cell clusters to cell type labels^68–70^.

### Differential expression analyses

For the smaller CRISPR *KO* lung screen, and the *OE* lung screen, pseudobulking and differential expression was performed using the Libra R package^52^, specifying edgeR’s LRT test. Specifically, cells were grouped by mouse (biological replicate), and each gRNA/*OE* construct was compared to the corresponding inert control construct. This was done separately for each cell type group, with the number of differentially expressed genes (FDR < 0.1) summarized in each case, ignoring instances with fewer than 20 cells per cell group. Only genes with minimum 5% total cell detection (per animal) were tested in all cases.

### Gene set enrichment analyses

Differentially expressed genes for select gRNAs/*OE* constructs were tested for enrichment against GO terms from MSigDB using the hypeR R package ^91^.

### Molecular feature effect analyses

Molecular feature sets curated (see section above) were used to score single cells based on their overall gene expression activity using pySCENIC’s AUCell implementation in Python. SCT-normalized counts were used to establish rankings, and molecular gene sets (lung / OA) were input as genesets in each case, yielding an AUCell activity score per cell per feature. The scores were then used as input to a linear regression model quantifying intervention vs corresponding control cell group effects (AUCell ∼ IntGrp). The corresponding regression β-coefficient and FDR-adjusted p values were used to determine the overall signed significance (sign(β) * −log10 *P_adj_*) for each intervention and feature.

### On-target gene expression fold change estimates

For visualizing the on-target effects in all screens taking cell number and percent target gene expression into account, we implemented a strategy that estimates fold-change even with cell number imbalance between groups. Briefly, for each intervention’s cells, we sampled the control cell group to match that of the intervention cell group size, running a Wilcoxon test for differential expression between the two groups for the corresponding target gene (*n*=100 iterations). We then determine the percent of iterations we observe a significant change in expression in the expected direction (downregulation for *KO* or *KD*, upregulation for *OE*, in the intervention relative to inert cells, Wilcoxon *P* < 0.05). Independently, for each target gene, we determined *k*=15 nearest-neighbor genes to each target gene matched on the basis of mean, percent and standard deviation in expression across all cells, and computed their corresponding fold-changes for the same cell groups being compared, generating what should be an empirical null around avg log FC=0.

### Intervention effect power analyses

For estimating sensitivity to detect on-target fold-changes, we took each barcode (gRNA, *OE* construct) and sampled the observed number of cells down to *n*=25 cells in steps of size 25. At each iteration, we computed the true positive rate (TPR) as the fraction of downsampled iterations that were significant at Wilcoxon *P* < 0.05 in the expected direction for intervention vs matched control downsampled cells. For estimating sensitivity to detect specific feature effects, we took each sampling iteration and re-ran the regression analysis of corresponding AUCell scores with respect to intervention vs inert groups. In either case, permuted cell groups (shuffled labels) were also compared and included in visualizations across all iterations.

### Human PCLS culture and siRNA transfection

Precision cut lung slices were generated as described previously^92^. Briefly, donor human lung tissue lobes were inflated with 1.5 - 2% low gelling agarose (Cat# A9414, Sigma-Aldrich) and allowed to solidify at 4 degrees for 30 min. 500 micron tissue slices were generated using vibratome (vibratome machine) at a cutting speed of 5 mm/sec. PCLS slices were maintained in DMEM/F12 culture medium with 1% FBS containing penicillin and streptomycin. For fibrosis induction, PCLS were treated with a fibrotic cocktail medium containing TGFB (5 ng/ml) (Cat# 781804, Biolegend), PDGFRB (10 ng/ml) (Cat#577306, Biolegend), TNFalpha (10 ng/ml) (Cat# 570102, Biolegend) and LPA (uM) (Cat# 10010093, Cayman) as described previously, medium was changed every other day.

For siRNA mediated knockdown studies, prevalidated siRNA against target genes were purchased from Dharmacon. PCLS were transfected with 500 ug of siRNA in 24 well format using Lipofectamine RNAiMAX reagent (Cat# 13778075, ThermoFisher) following manufacturers protocol. Total 3 doses of siRNA transfections were performed at day 7, 10, and 13 during the study period. siRNA mediated knockdown was assessed using real time quantitative PCR.

### RNA isolation and RT-qPCR

RNA isolation from PCLS was performed as described previously^93^. Two PCLS slices per donor from each condition were preserved in QIAzol Lysis Reagent (Cat# 79306, Qiagen) at −80 until processed. For RNA isolation, PCLS in QIAzol Lysis Reagent were homogenized using a beadbeater, 30 sec per cycle for a total of 4 cycles with cooling in between cycles for 1 min on ice. Homoginate was mixed with chloroform (0.1 ml of chloroform per 500 ul of QIAzol Lysis Reagent). Samples were then incubated for 5 min at room temperature and centrifuged for 15 min at 12 000 × g at 4 °C. To the 250 ul of aqueous phase, 1 ul (20 ug) of glycogen (Cat# 10901393001, Sigma-Aldrich), 250 ul isopropanol, and 120 ul of 5M of NaCl (final concentration is 1.2M) was added. Aqueous mix was vertex and incubated at −20 for 2 hours or overnight. Samples were then centrifuged for 10 min at 12000 × g at 4 °C to precipitate RNA, followed by washing the pellet three times with 70% ethanol at 12000x g for 5 min. The RNA pellet was air dried for 5 - 10 min, dissolved in 30 μl of RNase-free water, and incubated in a heat block set at 55 °C for 10 min. Total RNA content was measured using qubit and 300 ug of RNA was converted to cDNA using High-Capacity cDNA Reverse Transcription Kit (Cat# 4368814, ThermoFisher). qPCR assays were performed with the qTOWERiris Series (Analytikjena, California, USA). The relative quantities of mRNA for several genes were determined using iApplied Biosystems SYBR Green PCR Master Mix (Cat# 4309155, ThermoScientific). Target-gene transcripts in each sample were normalized to Glyceraldehyde-3-phosphate dehydrogenase (GAPDH) and expressed as a relative increase or decrease compared with control.

### Sircol assay

Sircol assay was performed following manufacturers protocol (Cat# S1000, Biocolor lifesciences). 2 PCLS slices per donor from each condition were digested using 0.1% (w/v) pepsin (Cat# 10108057001, Sigma-Aldrich) in 0.5 M acetic acid (1:20), maintained at pH 2.5 overnight. Following pepsin digestion, tissue digest was centrifuged at 3000x g for 10 min and supernatant was collected. To the 100 ul of tissue digest 1 ml of dye was added and placed on the shaker for 30 min. Collagen-dye complex was washed with 750 ul ice cold acid-salt wash reagent. Post wash the collagen bound dye was released using alkali reagent. Once the bound dye was dissolved, absorbance was measured along with standards using a spectrophotometer at 556 nm.

### Human knee joint cell and explant ex vivo cultures

Primary human osteoarthritic chondrocytes and synovial cells were isolated from tissue discards obtained from patients undergoing total knee arthroplasty (sourced from NDRI). Cells were transduced with an AAV2 vector encoding the transgene of interest under a CMV promoter at 100,000 MOI (5 × 10⁹ vg), with untransduced cells serving as controls. After 5 days in culture, conditioned media and papain-digested cells were assayed for sulfated GAG content using the dimethylmethylene blue (DMMB) assay. The dsDNA content was measured in papain-digested samples by Quant-iT Picogreen Assay (ThermoFisher Scientific) per manufacturer’s instructions for normalizing the GAG content.

### DMMB Assay

Sulfated glycosaminoglycan (sGAG) content was quantified using a DMMB assay. Briefly, explants or cell culture pellets were digested in papain buffer (125 µg/mL papain, 50 mM sodium phosphate, 2 mM EDTA, 2 mM L-cysteine, pH 6.5) at 60 °C for 16 h. Digested samples were centrifuged at 10,000 × g for 10 min, and supernatants were collected for analysis. DMMB dye reagent (16 mg/L DMMB, 3.04 g/L glycine, 1.6 g/L NaCl, pH 3.0) was added to each sample in a 1:10 ratio (v/v). Absorbance was measured immediately at 525 nm using a microplate reader. sGAG concentrations were calculated from a chondroitin-4-sulfate standard curve and normalized to DNA content measured by picogreen assay (Cat# P7589, Thermofisher Scientific) per manufacturer’s instructions.

### Statistical analysis

Functional data analysis was performed using GraphPad Prism 9 software (GraphPad Software, San Diego, CA, USA). For all experiments, data are shown as mean ± SEM, unless indicated otherwise. Comparisons between two groups were done using an unpaired two-tailed t-test, and comparisons of more than two groups were analyzed using one-way Brown-Forsythe ANOVA with Bonferroni correction. Figures and legends provide the number of independent experiments and the results of representative experiments.

### Data availability

GEO ID for single cell data generated as part of this study is GSE322491.

### Code availability

Code used for the processing and analyses of all the associated single cell RNAseq data generated in this study will be deposited to Github following data release.

## References

1. Dixit, A. et al. Perturb-seq: Dissecting molecular circuits with scalable single-cell RNA profiling of pooled genetic screens. Cell 167, 1853–1866.e17 (2016).

2. Adamson, B. et al. A multiplexed single-cell CRISPR screening platform enables systematic dissection of the unfolded protein response. Cell 167, 1867–1882.e21 (2016).

3. Borch Jensen, M. & Marblestone, A. In vivo pooled screening: A scalable tool to study the complexity of aging and age-related disease. Front. Aging 2, 714926 (2021).

4. Jin, X. et al. In vivo Perturb-Seq reveals neuronal and glial abnormalities associated with autism risk genes. Science 370, eaaz6063 (2020).

5. Santinha, A. J. et al. Transcriptional linkage analysis with in vivo AAV-Perturb-seq. Nature 622, 367–375 (2023).

6. Liu, S. J. et al. In vivo perturb-seq of cancer and microenvironment cells dissects oncologic drivers and radiotherapy responses in glioblastoma. Genomics (2023).

7. Zheng, X. et al. Massively parallel in vivo Perturb-seq reveals cell-type-specific transcriptional networks in cortical development. Cell 187, 3236–3248.e21 (2024).

8. Saunders, R. A. et al. A platform for multimodal in vivo pooled genetic screens reveals regulators of liver function. Genomics (2024).

9. Redente, E. F. et al. Persistent, progressive pulmonary fibrosis and epithelial remodeling in mice. Am. J. Respir. Cell Mol. Biol. 64, 669–676 (2021).

10. Izbicki, G., Segel, M. J., Christensen, T. G., Conner, M. W. & Breuer, R. Time course of bleomycin-induced lung fibrosis: Bleomycin-induced lung fibrosis. Int. J. Exp. Pathol. 83, 111–119 (2002).

11. Bordag, N. et al. Machine learning analysis of the bleomycin mouse model reveals the compartmental and temporal inflammatory pulmonary fingerprint. iScience 23, 101819 (2020).

12. Strunz, M. et al. Alveolar regeneration through a Krt8+ transitional stem cell state that persists in human lung fibrosis. Nat. Commun. 11, 3559 (2020).

13. Kobayashi, Y. et al. Persistence of a regeneration-associated, transitional alveolar epithelial cell state in pulmonary fibrosis. Nat. Cell Biol. 22, 934–946 (2020).

14. Bhatt, J., Ghigo, A. & Hirsch, E. PI3K/Akt in IPF: untangling fibrosis and charting therapies. Front. Immunol. 16, 1549277 (2025).

15. Sharma, P. et al. Molecular mechanisms and emerging therapeutics in pulmonary fibrosis: A recent update. Eur. J. Pharmacol. 1006, 178159 (2025).

16. Carriera, L. et al. Most promising emerging therapies for pulmonary fibrosis: Targeting novel pathways. Biomedicines 14, 154 (2026).

17. Sontake, V., Gajjala, P. R., Kasam, R. K. & Madala, S. K. New therapeutics based on emerging concepts in pulmonary fibrosis. Expert Opin. Ther. Targets 23, 69–81 (2019).

18. Hart, T. et al. Evaluation and design of genome-wide CRISPR/SpCas9 knockout screens. G3 (Bethesda) 7, 2719–2727 (2017).

19. Chen, C.-H. et al. Improved design and analysis of CRISPR knockout screens. Bioinformatics 34, 4095–4101 (2018).

20. Fernandez, I. E. & Eickelberg, O. The impact of TGF-β on lung fibrosis: from targeting to biomarkers. Proc. Am. Thorac. Soc. 9, 111–116 (2012).

21. Biernacka, A., Dobaczewski, M. & Frangogiannis, N. G. TGF-β signaling in fibrosis. Growth Factors 29, 196–202 (2011).

22. Hu, X., Li, J., Fu, M., Zhao, X. & Wang, W. The JAK/STAT signaling pathway: from bench to clinic. Signal Transduct. Target. Ther. 6, 402 (2021).

23. O’Shea, J. J. & Plenge, R. JAK and STAT signaling molecules in immunoregulation and immune-mediated disease. Immunity 36, 542–550 (2012).

24. Tsukui, T., Wolters, P. J. & Sheppard, D. Alveolar fibroblast lineage orchestrates lung inflammation and fibrosis. Nature 631, 627–634 (2024).

25. Aribindi, K., Liu, G. Y. & Albertson, T. E. Emerging pharmacological options in the treatment of idiopathic pulmonary fibrosis (IPF). Expert Rev. Clin. Pharmacol. 17, 817–835 (2024).

26. Bonella, F., Spagnolo, P. & Ryerson, C. Current and future treatment landscape for idiopathic pulmonary fibrosis. Drugs 83, 1581–1593 (2023).

27. Kampmann, M. CRISPRi and CRISPRa screens in mammalian cells for precision biology and medicine. ACS Chem. Biol. 13, 406–416 (2018).

28. Horlbeck, M. A. et al. Nucleosomes impede Cas9 access to DNA in vivo and in vitro. Elife 5, e12677 (2016).

29. Isaac, R. S. et al. Nucleosome breathing and remodeling constrain CRISPR-Cas9 function. Elife 5, e13450 (2016).

30. Modell, A. E., Lim, D., Nguyen, T. M., Sreekanth, V. & Choudhary, A. CRISPR-based therapeutics: current challenges and future applications. Trends Pharmacol. Sci. 43, 151–161 (2022).

31. Goyal, A. et al. Challenges of CRISPR/Cas9 applications for long non-coding RNA genes. Nucleic Acids Res. 45, e12 (2017).

32. Wu, Q. et al. Massively parallel characterization of CRISPR activator efficacy in human induced pluripotent stem cells and neurons. Mol. Cell 83, 1125–1139.e8 (2023).

33. Lalanne, J.-B. et al. Extensive length and homology dependent chimerism in pool-packaged AAV libraries. Genomics (2025).

34. Liu, T., Gonzalez De Los Santos, F., Hirsch, M., Wu, Z. & Phan, S. H. Noncanonical Wnt signaling promotes myofibroblast differentiation in pulmonary fibrosis. Am. J. Respir. Cell Mol. Biol. 65, 489–499 (2021).

35. Gottardi, C. J. & Königshoff, M. Considerations for targeting β-catenin signaling in fibrosis. Am. J. Respir. Crit. Care Med. 187, 566–568 (2013).

36. Wang, X., Zhu, H., Yang, X., Bi, Y. & Cui, S. Vasohibin attenuates bleomycin induced pulmonary fibrosis via inhibition of angiogenesis in mice. Pathology 42, 457–462 (2010).

37. Joannes, A. et al. FGF9 and FGF18 in idiopathic pulmonary fibrosis promote survival and migration and inhibit myofibroblast differentiation of human lung fibroblasts in vitro. Am. J. Physiol. Lung Cell. Mol. Physiol. 310, L615–29 (2016).

38. Yu, G. et al. Thyroid hormone inhibits lung fibrosis in mice by improving epithelial mitochondrial function. Nat. Med. 24, 39–49 (2018).

39. Giannandrea, M. & Parks, W. C. Diverse functions of matrix metalloproteinases during fibrosis. Dis. Model. Mech. 7, 193–203 (2014).

40. Pilling, D. & Gomer, R. H. Persistent lung inflammation and fibrosis in serum amyloid P component (APCs-/-) knockout mice. PLoS One 9, e93730 (2014).

41. Chen, J. et al. Enhanced detection of early pulmonary fibrosis disease using 68Ga-FAPI-LM3 PET. Mol. Pharm. 21, 3684–3692 (2024).

42. Lakatos, H. F. et al. The role of PPARs in lung fibrosis. PPAR Res. 2007, 71323 (2007).

43. Burgess, H. A. et al. PPARgamma agonists inhibit TGF-beta induced pulmonary myofibroblast differentiation and collagen production: implications for therapy of lung fibrosis. Am. J. Physiol. Lung Cell. Mol. Physiol. 288, L1146–53 (2005).

44. Wang, B. et al. The Kruppel-like factor KLF15 inhibits connective tissue growth factor (CTGF) expression in cardiac fibroblasts. J. Mol. Cell. Cardiol. 45, 193–197 (2008).

45. Wu, Z., Yang, H. & Colosi, P. Effect of genome size on AAV vector packaging. Mol. Ther. 18, 80–86 (2010).

46. Kugler, M. C. et al. Sonic Hedgehog signaling regulates myofibroblast function during alveolar septum formation in Murine postnatal lung. Am. J. Respir. Cell Mol. Biol. 57, 280–293 (2017).

47. Bolaños, A. L. et al. Role of Sonic Hedgehog in idiopathic pulmonary fibrosis. Am. J. Physiol. Lung Cell. Mol. Physiol. 303, L978–90 (2012).

48. Cao, H. et al. The Shh/Gli signaling cascade regulates myofibroblastic activation of lung-resident mesenchymal stem cells via the modulation of Wnt10a expression during pulmonary fibrogenesis. Lab. Invest. 100, 363–377 (2020).

49. Green, D. E., Kang, B.-Y., Murphy, T. C. & Hart, C. M. Peroxisome proliferator-activated receptor gamma (PPARγ) regulates thrombospondin-1 and Nox4 expression in hypoxia-induced human pulmonary artery smooth muscle cell proliferation. Pulm. Circ. 2, 483–491 (2012).

50. Guo, B. et al. Peroxisome proliferator-activated receptor-gamma ligands inhibit TGF-beta 1-induced fibronectin expression in glomerular mesangial cells. Diabetes 53, 200–208 (2004).

51. Lähnemann, D. et al. Eleven grand challenges in single-cell data science. Genome Biol. 21, 31 (2020).

52. Squair, J. W. et al. Confronting false discoveries in single-cell differential expression. Nature Communications 12, 5692 (2021).

53. Reyfman, P. A. et al. Single-cell transcriptomic analysis of human lung provides insights into the pathobiology of pulmonary fibrosis. Am. J. Respir. Crit. Care Med. 199, 1517–1536 (2019).

54. Kim, S. et al. Integrative phenotyping framework (iPF): integrative clustering of multiple omics data identifies novel lung disease subphenotypes. BMC Genomics 16, 924 (2015).

55. DePianto, D. J. et al. Heterogeneous gene expression signatures correspond to distinct lung pathologies and biomarkers of disease severity in idiopathic pulmonary fibrosis. Thorax 70, 48–56 (2015).

56. McDonough, J. E. et al. Gene correlation network analysis to identify regulatory factors in idiopathic pulmonary fibrosis. Thorax 74, 132–140 (2019).

57. Wang, Y. et al. Unsupervised gene expression analyses identify IPF-severity correlated signatures, associated genes and biomarkers. BMC Pulm. Med. 17, 133 (2017).

58. Ghandikota, S., Sharma, M., Ediga, H. H., Madala, S. K. & Jegga, A. G. Consensus gene co-expression network analysis identifies novel genes associated with severity of fibrotic lung disease. Int. J. Mol. Sci. 23, 5447 (2022).

59. Tsukui, T. et al. Collagen-producing lung cell atlas identifies multiple subsets with distinct localization and relevance to fibrosis. Nat. Commun. 11, 1920 (2020).

60. Habermann, A. C. et al. Single-cell RNA sequencing reveals profibrotic roles of distinct epithelial and mesenchymal lineages in pulmonary fibrosis. Sci. Adv. 6, eaba1972 (2020).

61. Nabhan, A. N., Brownfield, D. G., Harbury, P. B., Krasnow, M. A. & Desai, T. J. Single-cell Wnt signaling niches maintain stemness of alveolar type 2 cells. Science 359, 1118–1123 (2018).

62. Loeser, R. F., Goldring, S. R., Scanzello, C. R. & Goldring, M. B. Osteoarthritis: A Disease of the Joint as an Organ. Arthritis and Rheumatism 64, 1697 (2012).

63. Mathiessen, A. & Conaghan, P. G. Synovitis in osteoarthritis: current understanding with therapeutic implications. Arthritis research & therapy 19, (2017).

64. Malda, J. et al. Comparative study of depth-dependent characteristics of equine and human osteochondral tissue from the medial and lateral femoral condyles. Osteoarthritis and cartilage 20, (2012).

65. McIlwraith, C. W., Frisbie, D. D. & Kawcak, C. E. The horse as a model of naturally occurring osteoarthritis. Bone & joint research 1, (2012).

66. Sanchez-Lopez, E., Coras, R., Torres, A., Lane, N. E. & Guma, M. Synovial inflammation in osteoarthritis progression. Nature Reviews Rheumatology 18, 258–275 (2022).

67. Sellam, J. & Berenbaum, F. The role of synovitis in pathophysiology and clinical symptoms of osteoarthritis. Nature Reviews Rheumatology 6, 625–635 (2010).

68. Zhang, F. et al. Defining inflammatory cell states in rheumatoid arthritis joint synovial tissues by integrating single-cell transcriptomics and mass cytometry. Nature immunology 20, (2019).

69. Collins, F. L. et al. Taxonomy of fibroblasts and progenitors in the synovial joint at single-cell resolution. Annals of the Rheumatic Diseases 82, 428 (2022).

70. Micheroli, R. et al. Role of synovial fibroblast subsets across synovial pathotypes in rheumatoid arthritis: a deconvolution analysis. RMD open 8, (2022).

71. Tang, S. et al. Single-cell atlas of human infrapatellar fat pad and synovium implicates APOE signaling in osteoarthritis pathology. Sci. Transl. Med. 16, eadf4590 (2024).

72. He, W., Wang, M., Wang, Y., Wang, Q. & Luo, B. Plasma and Synovial Fluid CXCL12 Levels Are Correlated With Disease Severity in Patients With Knee Osteoarthritis. The Journal of arthroplasty 31, (2016).

73. Hadjadj, J. et al. Early-onset autoimmunity associated with SOCS1 haploinsufficiency. Nature communications 11, (2020).

74. Lee, P. Y. et al. Immune dysregulation and multisystem inflammatory syndrome in children (MIS-C) in individuals with haploinsufficiency of SOCS1. The Journal of allergy and clinical immunology 146, (2020).

75. Gui, T., He, B. S., Gan, Q. & Yang, C. Enhanced SOCS3 in osteoarthritis may limit both proliferation and inflammation. Biotechnic & Histochemistry 107–114 (2017).

76. Kaminski, N. et al. Global analysis of gene expression in pulmonary fibrosis reveals distinct programs regulating lung inflammation and fibrosis. Proc. Natl. Acad. Sci. U. S. A. 97, 1778–1783 (2000).

77. McDonough, J. E. et al. Transcriptional regulatory model of fibrosis progression in the human lung. JCI Insight 4, (2019).

78. Stein, Y. et al. Spatial and Single-cell Transcriptomics Reveal Programs Governing Fibroblastic Foci Fibroblasts. Genomics (2025).

79. Chin, D. et al. Genome-wide association study of Idiopathic Pulmonary Fibrosis susceptibility using clinically-curated European-ancestry datasets. Genetic and Genomic Medicine (2025).

80. Fingerlin, T. E. et al. Genome-wide association study identifies multiple susceptibility loci for pulmonary fibrosis. Nat. Genet. 45, 613–620 (2013).

81. Chiu, Y.-S. et al. The JAK inhibitor Tofacitinib inhibits structural damage in osteoarthritis by modulating JAK1/TNF-alpha/IL-6 signaling through Mir-149-5p. Bone 151, 116024 (2021).

82. Human osteoarthritic chondrocytes are impaired in matrix metalloproteinase-13 inhibition by IFN-γ due to reduced IFN-γ receptor levels. Osteoarthritis and Cartilage 17, 1049–1055 (2009).

83. Zheng, X., Thompson, P. C., White, C. M. & Jin, X. Massively parallel in vivo Perturb-seq screening. Nat. Protoc. 20, 1733–1767 (2025).

84. Shang, E., Wei, Y. & Roeder, K. Predicting the unseen: A diffusion-based debiasing framework for transcriptional response prediction at single-cell resolution. Proc. Natl. Acad. Sci. U. S. A. 122, e2525268122 (2025).

85. Lock, M. et al. Rapid, simple, and versatile manufacturing of recombinant adeno-associated viral vectors at scale. Hum. Gene Ther. 21, 1259–1271 (2010).

86. Konishi, S., Tata, A. & Tata, P. R. Defined conditions for long-term expansion of murine and human alveolar epithelial stem cells in three-dimensional cultures. STAR Protoc. 3, 101447 (2022).

87. Huang, L.-C. et al. scDemultiplex: An iterative beta-binomial model-based method for accurate demultiplexing with hashtag oligos. Comput Struct Biotechnol J 21, 4044–4055 (2023).

88. Korsunsky, I. et al. Fast, sensitive and accurate integration of single-cell data with Harmony. Nat Methods 16, 1289–1296 (2019).

89. Ma, F. & Pellegrini, M. ACTINN: automated identification of cell types in single cell RNA sequencing. Bioinformatics 36, 533–538 (2020).

90. Sikkema, L. et al. An integrated cell atlas of the lung in health and disease. Nat Med 29, 1563–1577 (2023).

91. Federico, A. & Monti, S. hypeR: an R package for geneset enrichment workflows. Bioinformatics 36, 1307–1308 (2019).

92. Gerckens, M. et al. Generation of human 3D lung tissue cultures (3D-LTCs) for disease modeling. J. Vis. Exp. e58437 (2019).

93. Langwiński, W. et al. An optimized QIAzol-based protocol for simultaneous miRNA, RNA, and protein isolation from precision-cut lung slices (PCLS). Respir. Res. 25, 422 (2024).

