## Supplementary figures & Table S1 for "A Multispecies, Modality-Agnostic Scalable In Vivo Mosaic Screening Platform for Therapeutic Target Discovery"

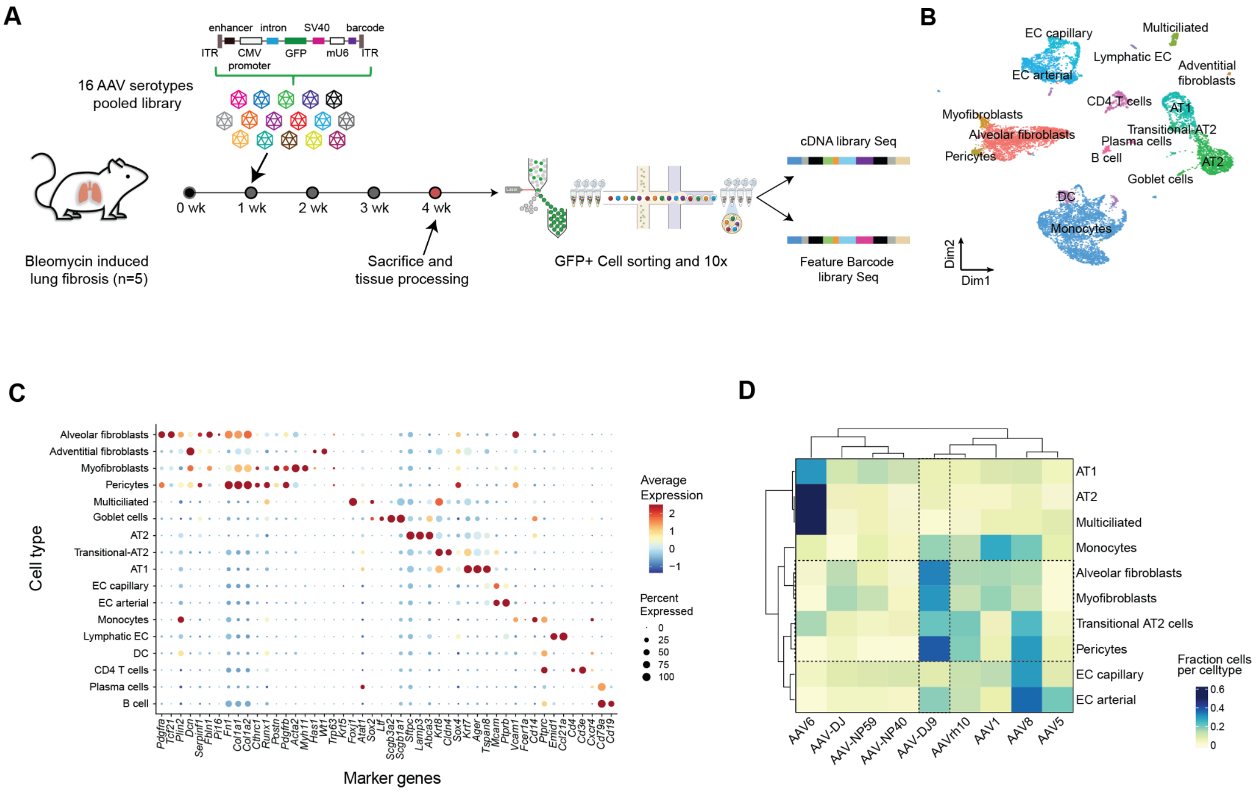


**Supplementary Fig S1. Identification of cell type-specific tropism of AAV through barcoded serotype screen in fibrotic lungs. A.** Schematic of fibrotic lung serotype screen experimental design. **B.** UMAP of cell types captured in fibrotic lung serotype screen **C.** Dot plot of cell type-specific gene expression markers. Dot size indicates the fraction of cells with detectable gene expression. Color scale indicates Z-score scaled average gene expression level. **D.** Heatmap showing cell type distribution per serotype barcode. Dotted box indicates cell types of interest and serotype used for subsequent mosaic screening.

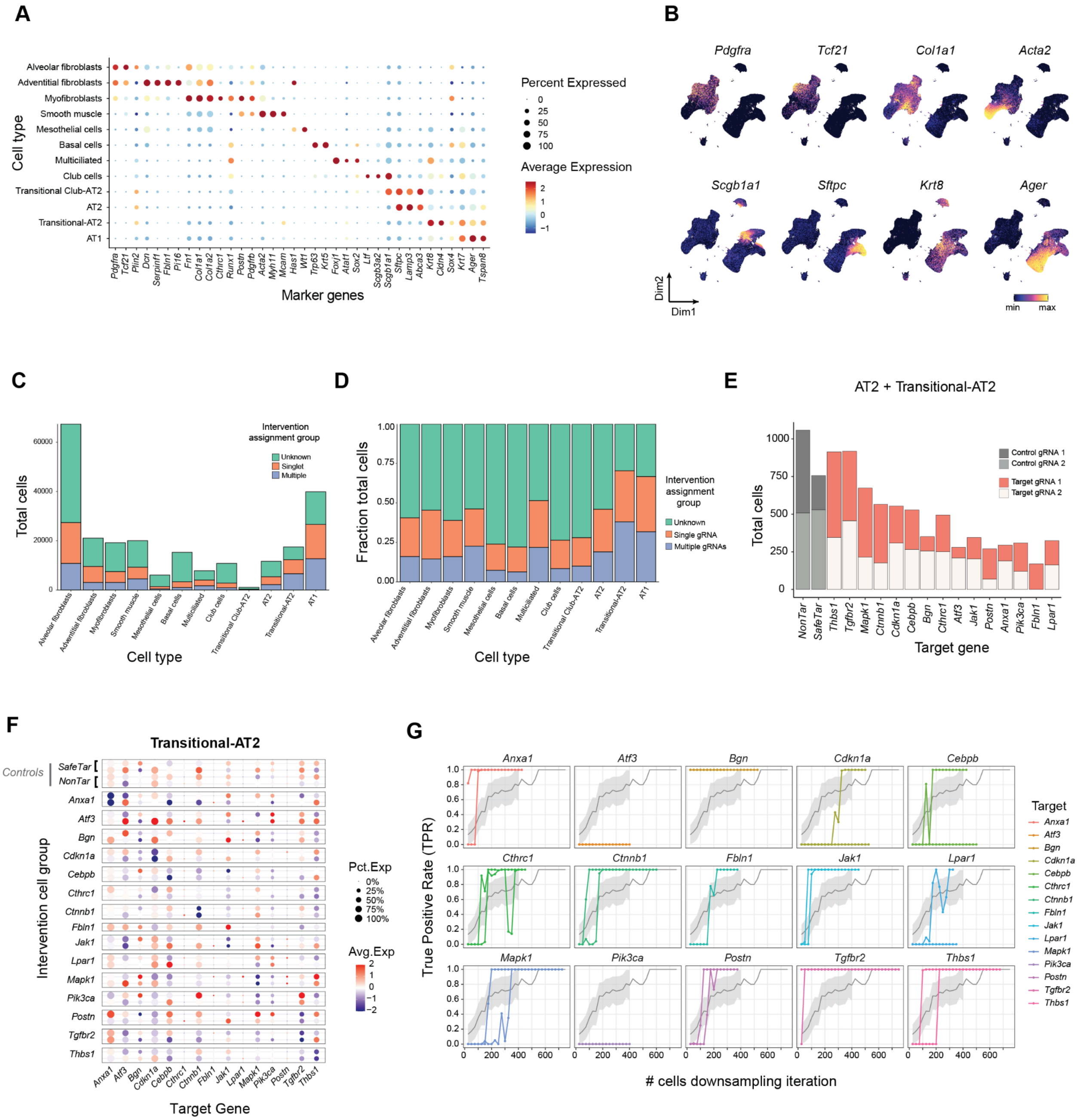
A
B
C
D
E
F
G
Cell type
Alveolar fibroblasts
Adventitial fibroblasts
Myofibroblasts
Smooth muscle
Mesothelial cells
Basal cells
Multiciliated
Club cells
Transitional Club-AT2
AT2
Transitional-AT2
AT1
Marker genes
Percent Expressed
Average Expression
Pdgfra
Tcf21
Col1a1
Acta2
Scgb1a1
Sftpc
Krt8
Ager
Dim2
Dim1
min
max
Total cells
Intervention assignment group
Unknown
Singlet
Multiple
Fraction total cells
Single gRNA
Multiple gRNAs
AT2 + Transitional-AT2
Control gRNA 1
Control gRNA 2
Target gRNA 1
Target gRNA 2
Target gene
Transitional-AT2
Controls
SafeTar
NonTar
Intervention cell group
Pct.Exp
Avg.Exp
Target Gene
True Positive Rate (TPR)
### cells downsampling iteration
Target
Anxa1
Atf3
Bgn
Cdkn1a
Cebpb
Cthrc1
Ctnnb1
Fbln1
Jak1
Lpar1
Mapk1
Pik3ca
Postn
Tgfbr2
Thbs1

**Supplementary Figure S2. Quality control and sensitivity analysis of the *in vivo* CRISPR KO mosaic screen. A.** Dot plot shows specific gene expression markers for annotated lung cell types in CRISPR KO screen. Dot size indicates the fraction of cells with detectable gene expression. Color scale indicates Z-score scaled average gene expression level. **B.** UMAP visualization of cell type-specific marker genes in fibrotic lungs. **C.** Cell count distribution of single, multiple and unknown gRNA assignments across cell types recovered from fibrotic lungs. **D.** Fractional distribution of single, multiple and unknown gRNA assignments across cell types. **E.** Number of single gRNA-assigned AT2 and transitional-AT2 cells recovered per target genes and controls. **F.** Dot plot showing scaled expression of library target genes in transitional-AT2 cells single-assigned to target specific gRNAs or controls (NonTar/SafeTar). Dot size indicates the fraction of cells with detectable gene expression. Color scale indicates Z-score scaled average gene expression. **G.** Scatter plot highlighting downsampled cell numbers (x-axis) versus true positive rate (TPR) computed as the fraction of total iterations ($n$=100) with significant knockdown (Wilcoxon $P$ < 0.05) for each gRNA relative to Non targeting control gRNA fibroblast cells. Gray: average with 95% confidence interval across all gRNAs-target pairs.

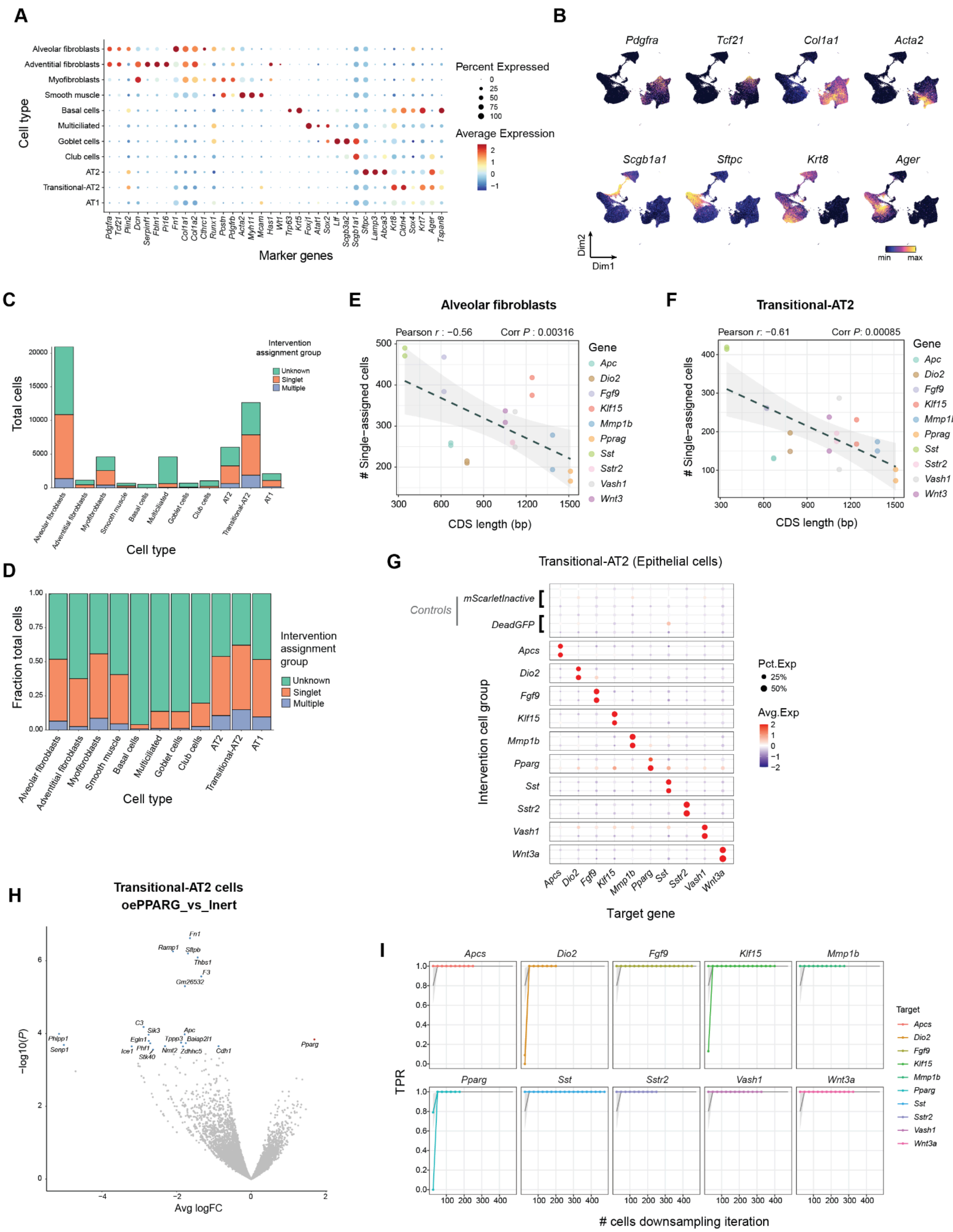
A
Cell type
Alveolar fibroblasts
Adventitial fibroblasts
Myofibroblasts
Smooth muscle
Basal cells
Multiciliated
Goblet cells
Club cells
AT2
Transitional-AT2
AT1
Marker genes
Percent Expressed
Average Expression
B
Pdgfra
Tcf21
Col1a1
Acta2
Scgb1a1
Sftpc
Krt8
Ager
Dim2
Dim1
min
max
C
Total cells
Intervention assignment group
Unknown
Singlet
Multiple
Cell type
D
Fraction total cells
Intervention assignment group
Unknown
Singlet
Multiple
Cell type
E
Alveolar fibroblasts
Pearson r : −0.56
Corr P : 0.00316
### Single−assigned cells
CDS length (bp)
Gene
Apc
Dio2
Fgf9
Klf15
Mmp1b
Pprag
Sst
Sstr2
Vash1
Wnt3
F
Transitional-AT2
Pearson r: −0.61
Corr P: 0.00085
### Single−assigned cells
CDS length (bp)
G
Transitional-AT2 (Epithelial cells)
Controls
mScarletInactive
DeadGFP
Intervention cell group
Target gene
Pct.Exp
25%
50%
Avg.Exp
H
Transitional-AT2 cells
oePPARG_vs_Inert
−log10(P)
Avg logFC
Pparg
I
Apcs
Dio2
Fgf9
Klf15
Mmp1b
Pparg
Sst
Sstr2
Vash1
Wnt3a
TPR
### cells downsampling iteration
Target

**Supplementary Figure S3. Quality control and assignment metrics for the in vivo *GOF* mosaic screen in fibrotic lungs. A.** Dot plot shows specific gene expression markers for annotated lung cell types in the *GOF* screen. Dot size indicates the fraction of cells with detectable gene expression. Color scale indicates Z-score scaled average gene expression level. **B.** UMAP visualization of cell type-specific marker genes annotated in fibrotic lungs. **C.** Cell count distribution of single, multiple and unknown barcode assignments across cell types recovered from fibrotic lungs. **D.** Fractional distribution of single, multiple, and unknown barcode assignments across cell types, with scatterplot highlighting inverse relationship between coding sequence length (CDS) and assignment rates in **E.** alveolar fibroblasts and **F.** transitional-AT2 cells**. G.** Dot plot showing scaled expression of library genes in transitional-AT2 cells receiving target *OE* construct and controls (deadGFP/mScarletInactive [mScarlet-I Y68A]). Dot size indicates the fraction of cells with detectable gene expression. Color scale indicates Z-score scaled average gene expression level. **H.** Volcano plots highlighting top differentially expressed genes by *Pparg* (Adj $P < 0.1$) in transitional-AT2 cells. **I.** Scatter plot highlighting downsampled cell numbers (x-axis) versus the true positive rate (TPR) computed as the fraction of total iterations ($n$=100) with significant over-expression (Wilcoxon $P < 0.05$) for each *OE* target relative to deadGFP control fibroblasts.

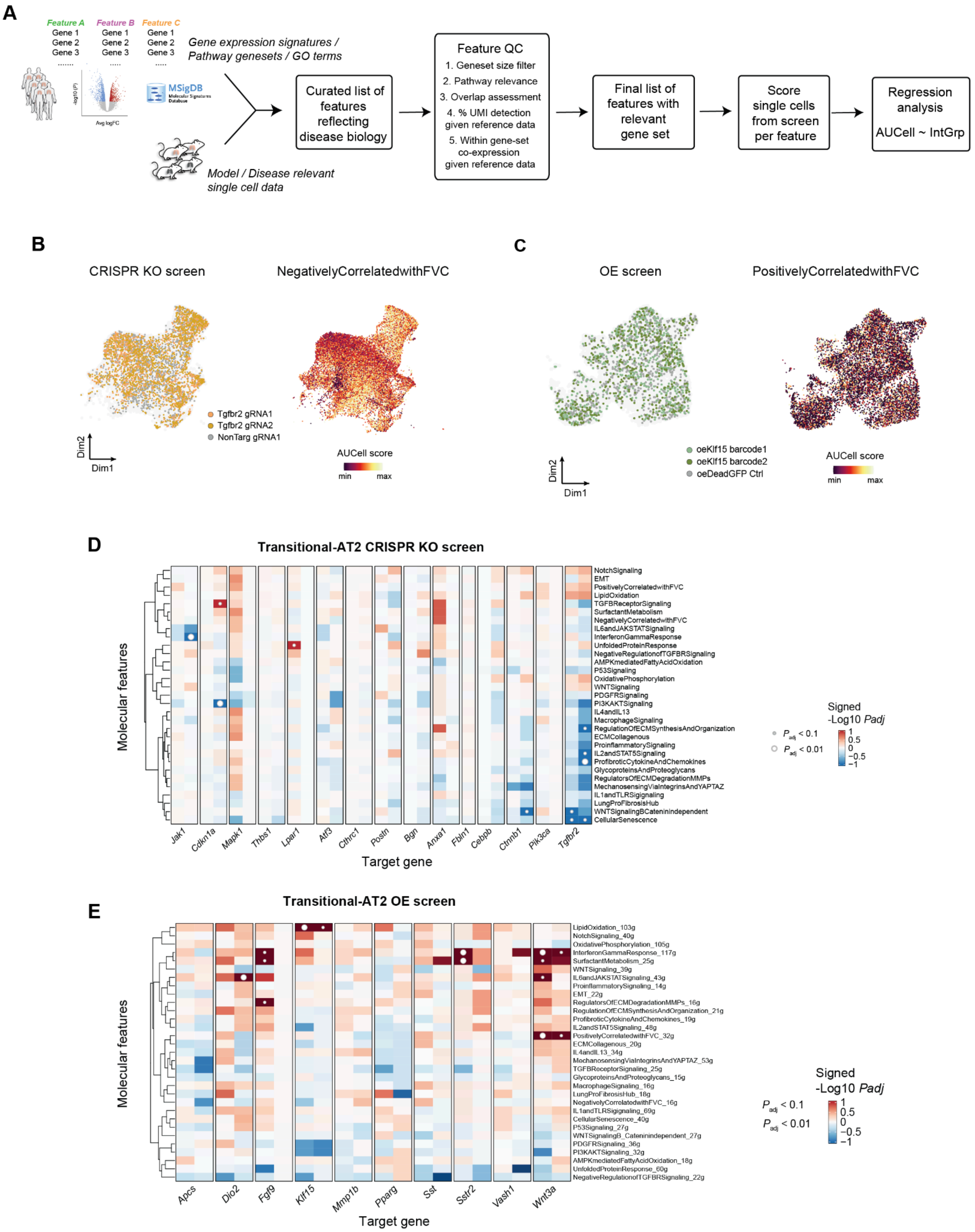
A
Feature A
Feature B
Feature C
Gene 1
Gene 2
Gene 3
Gene expression signatures / Pathway genesets / GO terms
MSigDB
Model / Disease relevant single cell data
Curated list of features reflecting disease biology
Feature QC
1. Geneset size filter
2. Pathway relevance
3. Overlap assessment
4. % UMI detection given reference data
5. Within gene-set co-expression given reference data
Final list of features with relevant gene set
Score single cells from screen per feature
Regression analysis
AUCell ~ IntGrp
B
CRISPR KO screen
NegativelyCorrelatedwithFVC
Tgfbr2 gRNA1
Tgfbr2 gRNA2
NonTarg gRNA1
Dim2
Dim1
AUCell score
min
max
C
OE screen
PositivelyCorrelatedwithFVC
oeKlf15 barcode1
oeKlf15 barcode2
oeDeadGFP Ctrl
Dim2
Dim1
AUCell score
min
max
D
Transitional-AT2 CRISPR KO screen
Molecular features
NotchSignaling
EMT
PositivelyCorrelatedwithFVC
LipidOxidation
TGFBReceptorSignaling
SurfactantMetabolism
NegativelyCorrelatedwithFVC
IL6andJAKSTATSignaling
InterferonGammaResponse
UnfoldedProteinResponse
NegativeRegulationofTGFBRSignaling
AMPKmediatedFattyAcidOxidation
P53Signaling
OxidativePhosphorylation
WNTSignaling
PDGFRSignaling
PI3KAKTSignaling
IL4andIL13
MacrophageSignaling
RegulationOfECMSynthesisAndOrganization
ECMCollagenous
ProinflammatorySignaling
IL2andSTAT5Signaling
ProfibroticCytokineAndChemokines
GlycoproteinsAndProteoglycans
RegulatorsOfECMDegradationMMPs
MechanosensingViaIntegrinsAndYAPTAZ
IL1andTLRSigignaling
LungProFibrosisHub
WNTSignalingBCateninindependent
CellularSenescence
Jak1
Cdkn1a
Mapk1
Thbs1
Lpar1
Atf3
Cthrc1
Postn
Bgn
Anxa1
Fbln1
Cebpb
Ctnnb1
Pik3ca
Tgfbr2
Target gene
Signed -Log10 Padj
Padj < 0.1
Padj < 0.01
1
0.5
0
−0.5
−1
E
Transitional-AT2 OE screen
Molecular features
LipidOxidation_103g
NotchSignaling_40g
OxidativePhosphorylation_105g
InterferonGammaResponse_117g
SurfactantMetabolism_25g
WNTSignaling_39g
IL6andJAKSTATSignaling_43g
ProinflammatorySignaling_14g
EMT_22g
RegulatorsOfECMDegradationMMPs_16g
RegulationOfECMSynthesisAndOrganization_21g
ProfibroticCytokineAndChemokines_19g
IL2andSTAT5Signaling_48g
PositivelyCorrelatedwithFVC_32g
ECMCollagenous_20g
IL4andIL13_34g
MechanosensingViaIntegrinsAndYAPTAZ_53g
TGFBReceptorSignaling_25g
GlycoproteinsAndProteoglycans_15g
MacrophageSignaling_16g
LungProFibrosisHub_18g
NegativelyCorrelatedwithFVC_16g
IL1andTLRSigignaling_69g
CellularSenescence_40g
P53Signaling_27g
WNTSignalingB_Cateninindependent_27g
PDGFRSignaling_36g
PI3KAKTSignaling_32g
AMPKmediatedFattyAcidOxidation_18g
UnfoldedProteinResponse_60g
NegativeRegulationofTGFBRSignaling_22g
Apcs
Dio2
Fgf9
Klf15
Mmp1b
Pparg
Sst
Sstr2
Vash1
Wnt3a
Target gene
Signed -Log10 Padj
Padj < 0.1
Padj < 0.01
1
0.5
0
−0.5
−1

**Supplementary Figure S4. Human molecular feature curation and quality control, and evaluation of transitional-AT2 cell type-specific effects A.** Schematic highlighting the feature gene set curation workflow, and corresponding utilization to quantify and assess screen intervention effects for physiologically-relevant features. **B.** UMAP of fibroblasts (CRISPR *KO* screen) colored by *Tgfbr2* and NonTar control gRNA assignments (left) or *Negatively Correlated with FVC* geneset AUCell score (right)**. C.** UMAP of fibroblasts (*GOF* screen) colored by *Klf15* and deadGFP control assignments (left) or *Positively Correlated with FVC* geneset AUCell score (right)**. D.** Heatmap of target gRNA vs NonTar control molecular feature effects. Color indicates signed -log10($P_{adj}$) from AUCell analysis of intervention vs control gRNA in transitional-AT2 cells. **E.** Heatmap of target *OE* vs deadGFP control molecular feature effects. Color indicates signed -log10($P_{adj}$) from AUCell scoring of *OE* intervention vs control in transitional-AT2 cells.

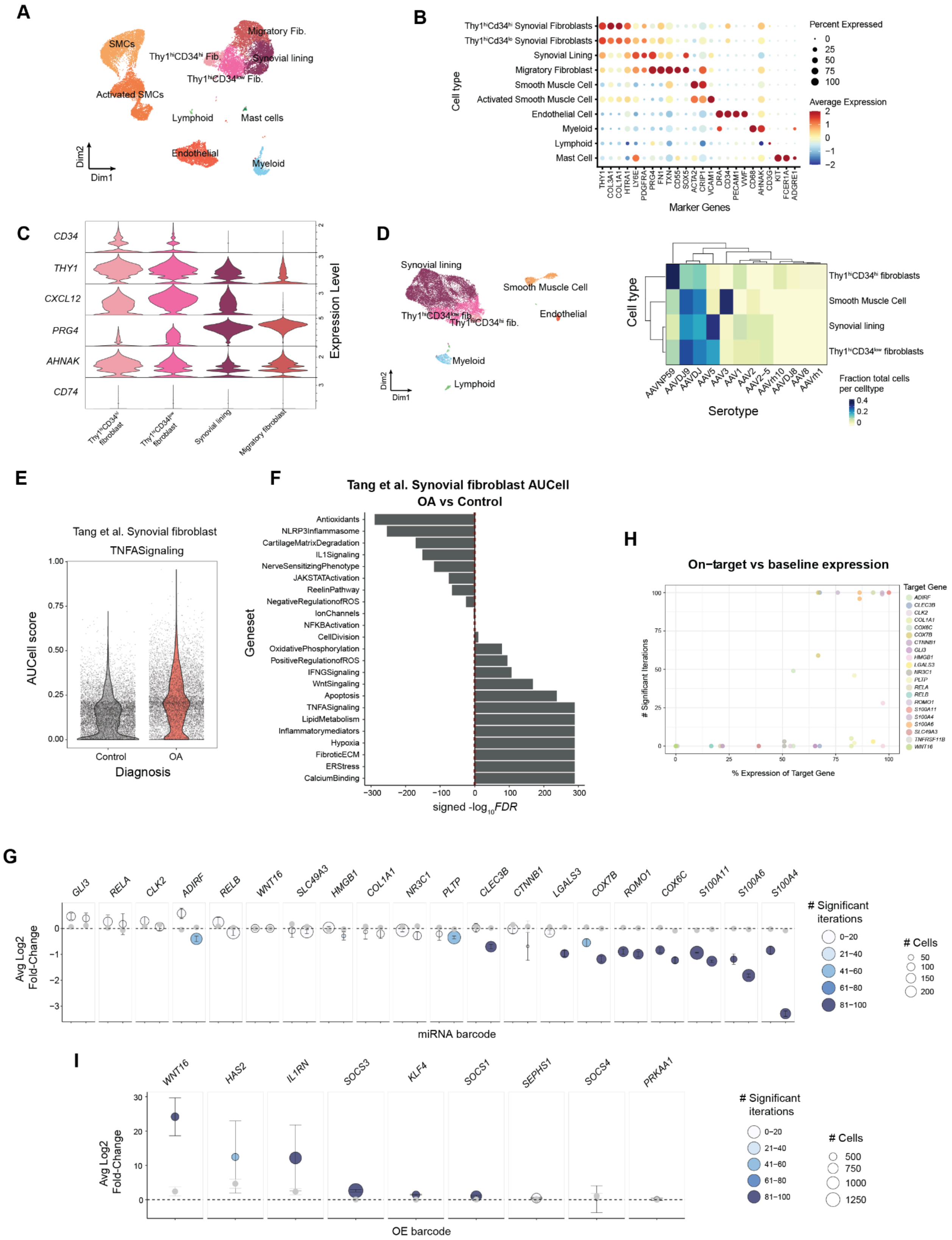
A
SMCs
Activated SMCs
Migratory Fib.
Synovial lining
Lymphoid
Mast cells
Endothelial
Myeloid
Dim2
Dim1
B
Thy1hiCd34hi Synovial Fibroblasts
Thy1hiCd34lo Synovial Fibroblasts
Synovial Lining
Migratory Fibroblast
Smooth Muscle Cell
Activated Smooth Muscle Cell
Endothelial Cell
Myeloid
Lymphoid
Mast Cell
Cell type
Marker Genes
Percent Expressed
Average Expression
C
CD34
THY1
CXCL12
PRG4
AHNAK
CD74
Expression Level
Synovial lining
Migratory fibroblast
D
Synovial lining
Smooth Muscle Cell
Endothelial
Myeloid
Lymphoid
Cell type
Serotype
Thy1hiCD34hi fibroblasts
Smooth Muscle Cell
Synovial lining
Thy1hiCD34low fibroblasts
Fraction total cells per celltype
E
Tang et al. Synovial fibroblast
TNFASignaling
AUCell score
Control
OA
Diagnosis
F
Tang et al. Synovial fibroblast AUCell
OA vs Control
Antioxidants
NLRP3Inflammasome
CartilageMatrixDegradation
IL1Signaling
NerveSensitizingPhenotype
JAKSTATActivation
ReelinPathway
NegativeRegulationofROS
IonChannels
NFKBActivation
CellDivision
OxidativePhosphorylation
PositiveRegulationofROS
IFNGSignaling
WntSingaling
Apoptosis
TNFASignaling
LipidMetabolism
Inflammatorymediators
Hypoxia
FibroticECM
ERStress
CalciumBinding
Geneset
signed -log10FDR
H
On−target vs baseline expression
Target Gene
### Significant Iterations
% Expression of Target Gene
G
GLI3
RELA
CLK2
ADIRF
RELB
WNT16
SLC49A3
HMGB1
COL1A1
NR3C1
PLTP
CLEC3B
CTNNB1
LGALS3
COX7B
ROMO1
COX6C
S100A11
S100A6
S100A4
Avg Log2 Fold-Change
miRNA barcode
### Significant iterations
### Cells
I
WNT16
HAS2
IL1RN
SOCS3
KLF4
SOCS1
SEPHS1
SOCS4
PRKAA1
OE barcode

**Supplementary Figure S5. Single-cell profiling of equine synovial cell types, serotype tropism, and on-target effects. A.** UMAP of horse synovial cells (n=2) from naive OA horse fetlock joints. **B.** Dot plot of marker gene expression per annotated cell type group for horse synovial tissue. Dot size indicates the fraction of cells with detectable gene expression. Color scale indicates Z-score scaled average gene expression level. **C.** Violin plot of synovial marker gene expression comparing the different synovial cell subtypes. **D**. (Left) UMAP of cells with annotated cell types shown in text. (Right) Serotype screen performed using single cell sequencing of horse synovial samples (*n*=2 horses, *n*=4 joints). Heatmap showing the fraction of total single-assigned cells for a given serotype barcode assigned to each cell annotation group. **E.** Violin plot of AUCell scores for the *TNFASignaling* feature corresponding to synovial fibroblasts in healthy and OA human synovial fibroblast samples as described in Tang et al. **F**. Waterfall plot of OA vs Ctrl differential AUCell scores (signed -log10 Wilcoxon FDR) for OA disease features in synovial fibroblasts from published human scRNA-seq data. **G**. Scatter plot of average log2 fold-change for each miRNA barcode vs scrambled control-receiving synovial cells. Dot size represents the number of cells, while the number of significant (Wilcoxon $P < 0.05$, log2FC < 0) iterations is determined from $n=100$ sampling iterations of equal cell numbers between groups. Gray points and error bars indicate null hypothesis, as mean +/- s.e. Avg log2FC between the corresponding two cell groups, but for randomly matched nearest-neighbor genes to the target gene. **H.** Spearman correlation between expression of target gene in miRNA screen versus significant iteration of KD effect for corresponding target. **I.** Same as in G., but for horse synovial *OE* screen, and with significant iterations computed for target gene upregulation (Wilcoxon $P < 0.05$, log2FC > 0).

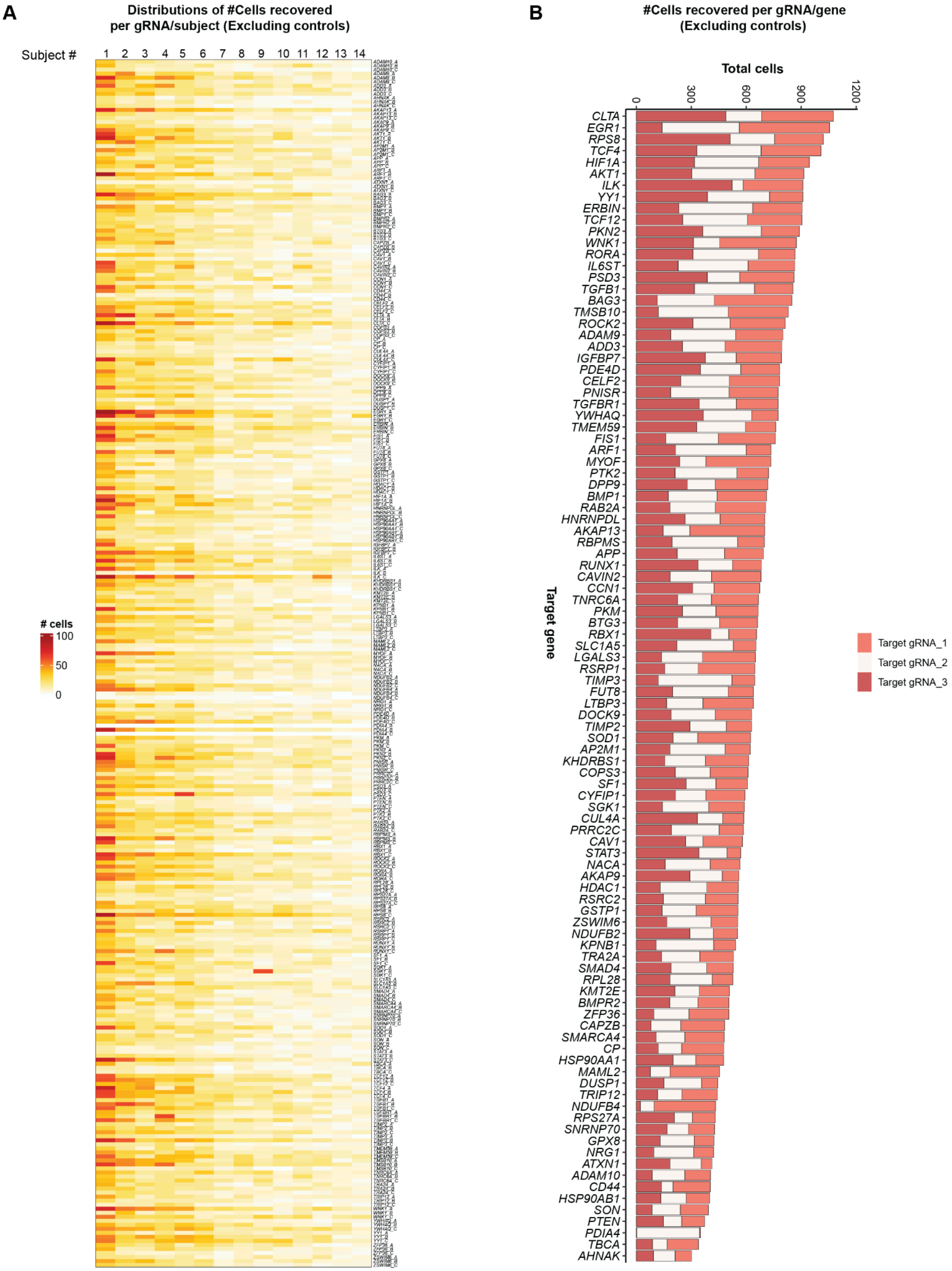
A
Distributions of #Cells recovered
per gRNA/subject (Excluding controls)
Subject #
1 2 3 4 5 6 7 8 9 10 11 12 13 14
### cells
100
50
0
B
#Cells recovered per gRNA/gene
(Excluding controls)
Total cells
0
300
600
900
1200
Target gene
CLTA
EGR1
RPS8
TCF4
HIF1A
AKT1
ILK
YY1
ERBIN
TCF12
PKN2
WNK1
RORA
IL6ST
PSD3
TGFB1
BAG3
TMSB10
ROCK2
ADAM9
ADD3
IGFBP7
PDE4D
CELF2
PNISR
TGFBR1
YWHAQ
TMEM59
FIS1
ARF1
MYOF
PTK2
DPP9
BMP1
RAB2A
HNRNPDL
AKAP13
RBPMS
APP
RUNX1
CAVIN2
CCN1
TNRC6A
PKM
BTG3
RBX1
SLC1A5
LGALS3
RSRP1
TIMP3
FUT8
LTBP3
DOCK9
TIMP2
SOD1
AP2M1
KHDRBS1
COPS3
SF1
CYFIP1
SGK1
CUL4A
PRRC2C
CAV1
STAT3
NACA
AKAP9
HDAC1
RSRC2
GSTP1
ZSWIM6
NDUFB2
KPNB1
TRA2A
SMAD4
RPL28
KMT2E
BMPR2
ZFP36
CAPZB
SMARCA4
CP
HSP90AA1
MAML2
DUSP1
TRIP12
NDUFB4
RPS27A
SNRNP70
GPX8
NRG1
ATXN1
ADAM10
CD44
HSP90AB1
SON
PTEN
PDIA4
TBCA
AHNAK
Target gRNA_1
Target gRNA_2
Target gRNA_3

**Supplementary Figure S6. Distribution and recovery of gRNAs assigned cells across experimental subjects per target genes in fibroblasts. A.** Heatmap of distribution of the number of cells recovered for each individual gRNA across 14 experimental subjects. The color gradient, ranging from light yellow to dark red, represents the absolute cell count per gRNA/subject, demonstrating consistent coverage across the cohort. **B.** Bar chart showing cell counts per gRNA per target gene captured. The bars are subdivided into three segments (Target gRNA_1, Target gRNA_2, Target gRNA_3), representing the recovery of the specific gRNAs assigned to a single gene.

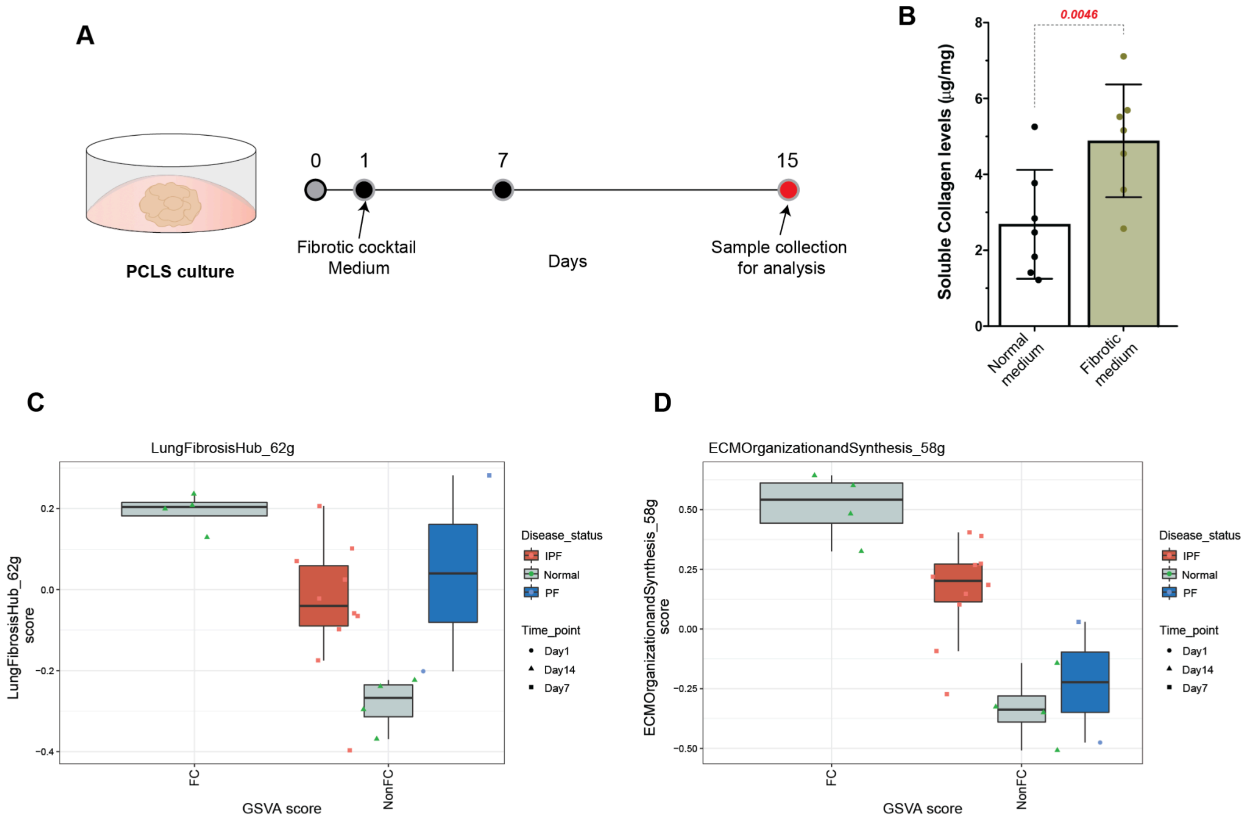


**Supplementary Figure S7: Human precision cut lung slices treated with fibrotic cocktail models fibrosis phenotype. A.** Schematic of PCLS culture workflow. Donor tissue slices were cultured in a fibrotic cocktail medium from Day 1 to Day 14, with samples collected for analysis at day 7 and day 14. **B.** Bar graph showing elevated soluble collagen levels (µg/mg) in PCLS treated with fibrotic medium compared to normal medium. **C-D.** Boxplots showing Gene Set Variation Analysis (GSVA) scores for the *Lung Pro-Fibrosis Hub* (C) and *ECM Organization and Synthesis* (D) features across different time points (Day 1, 7, 14), comparing fibrotic cocktail to untreated tissues from idiopathic pulmonary fibrosis, pulmonary fibrosis, and healthy control.

#### Table S1: Curated gene sets per molecular features for PF screens

| PF Molecular Feature | Genes |
|---|---|
| PositivelyCorrelatedwith FVC | *ADRB2,AFF3,NINJ2,FRY,ARHGAP31,ARHGEF26,ANKS1A,CCBE1,NCKAP5,EPB41L5,ANXA3,WNT3A,EMP2,FZD5,SPRYD7,SLC44A2,KCNMB4,OLFML2A,ECHDC3,SEMA3B,LAMA3,PCYOX1,RNF144B,HYAL1,CDH13,CTNND2,DPP6,GRIA1,DENND3,RTKN2,AGER,WNT7A,MYRF* |
| NegativelyCorrelatedwithFVC | *IGF1,LTBP1,SULF1,COL15A1,SERPINF1,SFRP2,PDIA4,COL14A1,COL18A1,GPX8,COL3A1,STEAP2,STEAP1,TTC39C,DCLK1,ITGA7* |
| LungProFibrosisHub | *ELN,FAM13A,FGF2,MECP2,MT2A,CCN2,TIMP1,ELMOD2,MMP2,HGF,ATP11A,CEBPB,PARN,DPP9,HMOX1,SKIL,STN1,PTX3,NFE2L2* |
| ProfibroticCytokineAndChemokines | *TGFB2,TGFB1,CSF2,CXCL8,CCL11,IL13,TNF,CXCL2,MMP9,IL4,IL6,IL5,IL1B,CCL5,CCL4,SPP1,CCL3,CCL2,PDGFB,PDGFA* |
| ECMCollagenous | *COL1A1,COL5A1,COL3A1,COL7A1,COL21A1,COL24A1,COL27A1,COL28A1,COL4A1,COL4A2,COL4A4,COL4A5,COL4A6,COL5A2,COL5A3,COL6A1,COL6A2,COL6A3,COL6A5,COL6A6,COL8A1* |
| GlycoproteinsAndProteoglycans | *DCN,ELN,LAMC2,LAMB1,FN1,NID1,LAMA5,LAMC1,HSPG2,FBN1,FBLN1,HAS1,HAS2,SULF1,VCAN* |
| RegulatorsOfECMDegradation MMPs | *BSG,MMP14,MMP11,MMP15,FURIN,TCF20,MMP2,CTSK,MMP3,MMP19,MMP13,MMP9,MMP16,ADAM22,MMP10,MMP17* |
| RegulationOfECMSynthesisAndOrganization | *CD44,PSEN1,FURIN,SCUBE1,NCSTN,BMP1,PHYKPL,SFRP2,P4HA3,COLGALT1,CRTAP,PCOLCE,PCOLCE2,PLOD1,PLOD2,PLOD3,P4HB,PPIB,P3H1,P3H3,SERPINH1* |
| ProinflammatorySignaling | *IL1RN,CXCL9,CSF1,IFNA1,EPO,CXCL1,IL27,CXCL5,TNFSF13B,IL36B,IL10,IL15,MMP1,IL1A,IFNG,LTA,IL36RN,CCL18,CCL17* |
| IL1andTLRSigignaling | *CHUK,ERC1,IKBKG,IL1R2,IRAK4,MAP3K3,PIK3CA,PIK3R1,PRKCI,PRKCZ,SQSTM1,TAB2,TICAM2,UBE2N,UBE2V1,IKBKB,IL1A,IL1B,IL1R1,IL1RAP,IL1RN,IRAK1,IRAK3,JUN,MAP2K6,MAP3K7,MAPK8,MYD88,NFKB1,RELA,TAB1,TOLLIP,TRAF6,ECSIT,IFNA1,IFNB1,IL6,IRAK2,MAP2K3,MAP3K1,MAP3K14,MAPK14,NFKBIA,TGFB1,TGFB2,TGFB3,TNF,ATF2,CCL2,HSPB2,MAP2K1,MAP2K2,MAP2K4,MAP2K7,MAP3K2,MAPK1,MAPK3,MAPK9,MAPKAPK2,NFKBIB,PELI1,PELI2,PIK3R2,PLCG1,PTPN11,REL,TAB3,CTSG,GSDMD,IL18,NFKB2* |
| IL2andSTAT5Signaling | *AHR,BATF,BATF3,BCL2,BCL2L1,CCND2,CCND3,CCNE1,CD44,CD83,CD86,CDC6,CISH,CSF2,CTLA4,CXCL10,EOMES,FURIN,GADD45B,GPR83,HK2,ICOS,IFNGR1,IKZF2,IKZF4,IL10,IL13,IL1RL1,IL2RA,IL2RB,IL4R,IRF4,ITGAE,LIF,MYC,NFIL3,NRP1,NT5E,ODC1,PIM1,SELL,SLC1A5,SLC2A3,SOCS1,SOCS2,TNFRSF18,TNFRSF4,TNFRSF8,TNFRSF9,TRAF1* |
| IL6andJAKSTATSignaling | *IL6,ACVR1B,ACVRL1,BAK1,CBL,CD44,CD9,CNTFR,CRLF2,CSF1,CSF2RA,CXCL1,CXCL10,CXCL9,FAS,GRB2,HAX1,HMOX1,IFNAR1,IFNGR1,IL10RB,JAK2,PIAS4,STAT1,STAT3,STAT5A,STAT5B,STAT6,PTPN11,GRB2,IL2RG,MYD88,TNFRSF1A,STAT4,TYK2,JAK1,IL6R,IL6ST,SOCS3,PIM1,PTGS2,OSMR,SOCS1,PTPN1,PTPN2* |
| IL4andIL13 | *IL4,IL13,IL4R,IL13RA1,IL13RA2,JAK3,PIK3R1,GATA3,IRF4,BATF,ALOX15,ALOX5,ARG1,BCL2,BCL2L1,CCL2,CCL11,CCL22,CD36,COL1A2,FCER2,FN1,F13A1,FSCN1,IGHE,IGHG1,IGHG4,IL10,ITGAM,LAMA5,LCN2,MAOA,MMP1/2/3/9,MUC1,RHOU,TIMP1,VCAM1,VEGFA,ZEB1/TWIST1,ITGAX* |
| InterferonGammaResponse | *ADAR,ARID5B,ARL4A,AUTS2,B2M,BST2,BTG1,CASP4,CASP7,CD40,CD74,CDKN1A,CFB,CFH,CMKLR1,DDX58,DDX60,EIF2AK2,EIF4E3,FAS,FGL2,GBP6,GCH1,ICAM1,IFI27,IFI30,IFI35,IFIH1,IFIT2,IFIT3,IFITM2,IFITM3,IFNAR2,IL15,IRF1,IRF2,IRF7,IRF8,IRF9,ISG15,ISG20,ISOC1,JAK2,LAP3,LATS2,LGALS3BP,LY6E,LYSMD2,MVP,NAMPT,NCOA3,NFKB1,NFKBIA,NMI,NOD1,NUP93,OGFR,P2RY14,PARP12,PARP14,PDE4B,PELI1,PFKP,PIM1,PLA2G4A,PLSCR1,PML,PNP,PNPT1,PSMA2,PSMA3,PSMB10,PSMB2,PSMB8,PSMB9,PSME1,PSME2,PTGS2,RAPGEF6,RBCK1,RIPK1,RIPK2,RNF213,RNF31,RTP4,SAMD9L,SAMHD1,SERPING1,SLC25A28,SOCS1,SOCS3,SOD2,SP110,SPPL2A,SRI,SSPN,ST3GAL5,TAP1,TAPBP,TDRD7,TNFAIP2,TNFAIP3,TNFAIP6,TOR1B,TRAFD1,TRIM25,TRIM26,TXNIP,UBE2L6,USP18,VAMP5,VAMP8,VCAM1,WARS,XAF1,ZBP1,ZNFX1* |
| MacrophageSignaling | *CSF1,CCL2,CXCL16,C3,PDGFRA,TGFBR2,IL1R1,EGFR,STAT1,CCN2,CXCL12,STAT3,STAT6,NFE2L2,IL6,IL12A* |
| CellularSenescence | *CDKN1B,KAT5,SCMH1,KDM6B,RING1,EED,TERF2,IL1A,H2AJ,TFDP1,MAPKAPK3,TFDP2,TP53,RNF2,FZR1,RBBP4,E2F1,E2F2,E2F3,RBBP7,CBX8,CDKN2D,SUZ12,CBX6,PHC2,CDKN2B,PHC1,CDKN2C,CBX4,CDKN2A,IFNB1,CBX2,HMGA1,HMGA2,PHC3,CDK6,CDK4,CDK2,MDM2,ATM,MDM4,EZH2* |
| UnfoldedProteinResponse | *HSPA5,CALR,EIF4A1,ATF3,PDIA6,NPM1,EEF2,HSP90B1,YWHAZ,RPS14,CEBPB,LMNA,SPCS1,DDIT4,HSPA9,ATF4,LSM4,BANF1,HDGF,BAG3,EIF2S1,HERPUD1,EIF4G1,NHP2,XBP1,EIF4A3,SEC11A,NOLC1,EIF2S2,FUS,DNAJB9,DNAJC3,NOP56,DNAJB11,EIF4A2,PSAT1,KIF5B,HYOU1,SERP1,MYDGF,EIF4E,EIF4EBP1,SSR1,IMP3,CKS1B,ACADVL,TPP1,SDAD1,YIF1A,SEC31A,NOP14,SPCS3,NFYB,DKC1,EXOSC8,GSK3A,DCTN1,ASNS,EXOSC5,ATP6V0D1* |

| | |
|---|---|
| OxidativePhosphorylation | *COX8A,COX6C,COX4I1,NDUFA4,COX7A2,NDUFA7,UQCRB,COX6B1,COX6A1,COX5A,UQCRH,SLC25A5,UQCR11,LDHA,COX7B,ATP6V1G1,UQCRQ,UQCR10,ATP6V0C,NDUFC2,COX5B,COX7C,NDUFC1,MDH1,NDUFA6,ETFB,NDUFB8,CYB5A,OAT,LDHB,NDUFB5,NDUFA2,NDUFA3,SLC25A3,NDUFB6,IDH2,GPX4,HSD17B10,ATP1B1,NDUFB4,CYCS,NDUFAB1,NDUFA1,NDUFB2,MGST3,MDH2,MPC1,CYC1,NDUFB3,NDUFA8,PRDX3,NDUFA5,NDUFS6,ECHS1,NDUFV2,NDUFS7,COX7A2L,UQCRFS1,SDHB,TIMM13,ATP6V1F,NDUFS8,UQCRC1,TIMM8B,ISCU,CYB5R3,NDUFS3,TOMM22,SDHC,COX17,VDAC1,MRPL34,NDUFB7,ECI1,SUCLG1,BAX,FDX1,SDHD,GRPEL1,ATP6V0B,VDAC2,ACAA2,TIMM17A,ECH1,UQCRC2,PHB2,ETFA,POLR2F,NDUFS2,SLC25A11,VDAC3,NDUFV1,MRPS12,ALAS1,NDUFA9,ATP6V1E1,ACO2,NDUFS4,ACAT1,IDH3A,ATP6V1D,IDH3B,MRPL11,RETSAT,MRPL15* |
| P53Signaling | *STEAP3,CD82,EI24,PPM1D,RCHY1,PIDD1,CCND3,CCND2,ZMAT3,SESN1,PERP,SFN,APAF1,GADD45A,CDKN2A,SERPINB5,DDB2,CDK6,TRP53,CDK2,CCNG1,MDM2,FAS,BAX,ATM,MDM4,TRP53,CDK4,TP53* |
| TGFBReceptorSignaling | *TGFB1,INHBA,FURIN,LTBP1,ITGB1,ITGB6,TGFBR1,TGFBR2,TGFBR3,SMAD1,SMAD2,SMAD3,SMAD4,SMAD5,RHOA,ZFYVE9,ZFYVE16,USP15,UCHL5,RNF111,EP300,SNW1,SP1,E2F4,TFDP1* |
| DownRegulationofTGFBRSignaling | *PPP1R15A,SMURF2,SMURF1,NEDD4L,MTMR4,SMAD7,PPP1CA,PPP1CB,PPP1CC,XPO1,BAMBI,STRAP,PMEPA1,STUB1,SMAD6* |
| MechanosensingViaIntegrinsAndYAPTAZ | ACTA2,ACTB,ACTG1,CDH1,CTNNA1,CTNNB1,ITGB1,ITGB5,ITGB8,LATS1,LIMD1,NF2,PAK1,PAK2,PAK4,ITGB1,ITGB2,ITGB3,ITGB4,ITGB5,ITGB6,ITGB7,ITGB8,ITGBL1,LYN,PLP1,TSPAN32,COL1A2,EMILIN1,ITGA1,ITGA10,ITGA11,ITGA2,ITGA2B,ITGA3,ITGA4,ITGA5,ITGA6,ITGA7,ITGA8,ITGA9,ITGAD,ITGAE,ITGAL,ITGAM,ITGAV,ITGAX,TEAD1,WWTR1,MAP4K3,MAP4K4,YWHAQ,SRC,STK3,YAP1,MAP4K5 |
| PI3KAKTSignaling | CDKN1A,CDKN1B,PRKAA2,YWHAB,PTEN,PIK3R3,FASLG,IL2RG,EGFR,HSP90B1,GNGT1,RPTOR,FGF6,CCND2,CCND1,PPP2R1B,MYC,AKT1,THEM4,MAPK1,ITGAV,RAC1,HRAS,EIF4E,FGF22,TSC2,PRKCA,NGF,FGF17,IL4,CDK4,CDK2,GRB2,RAF1 |
| NotchSignaling | DLL1,DTX1,DTX2,DTX4,HES1,JAG1,KAT2A,LFNG,NOTCH1,NOTCH2,NOTCH3,PSEN2,MAML2,MAML1,HDAC1,NOTCH4,CUL1,ARRB1,PSEN1,RFNG,RBPJ,APH1A,DLL4,NCSTN,APH1B,CCND1,ST3GAL6,HES5,SKP1,PTCRA,PSENEN,JAG2,CREBBP,RBX1,NCOR2,KAT2B,HEYL,SNW1,MFNG,NUMB,MAML3 |
| PDGFRSignaling | PDGFRB,PDGFRA,CRK,RAPGEF1,CRKL,ITGAV,SHB,CAV1,SHF,AKT1,CAV3,GSK3B,LRP1,PRKCA,PRKCB,RAC1,CDK1,PPP2CA,HRAS,CDC25C,CCND1,CCND2,SKP2,RUNX2,CUL1,MAP2K1,MAP2K4,MAP3K1,MAP3K7,MAPK1,MAPK3,MAPK8,MAPK8IP1,MAPK9,RAF1,RASA1 |
| WNTSignaling | FRAT1,FRAT2,RUVBL1,AKT1,EP300,SOX9,BTRC,CSNK2A1,CTNNBIP1,CSNK2A2,AXIN1,CSNK1E,DKK1,SFRP1,SFRP2,CSNK2B,TCF7,CUL1,LRP5,CXXC4,LRP6,PPP2CA,WNT6,WNT2,WNT3,WNT3A,CSNK1A1,CAV1,WNT7A,BCL9,TCF4,TCF3,SKP2,CCND2,HEY1,HEY2,GNAI1,NKD1,PPARD |
| WNTSignalingB_Catenindependent | ARRB2,MYC,RAC1,MAP3K7,WNT5B,PRKCB,WNT5A,PRKCA,AXIN2,RHOA,DAAM1,ROR2,LEF1,CAMK2A,NLK,WNT1,WNT4,FZD1,FZD2,FZD5,FZD4,FZD7,FZD6,FZD8,VANGL2,CTNNB1,FZD9 |
| SurfactantMetabolism | ABCA3,ADA2,ADGRF5,ADORA2A,ADORA2B,ADRA2A,ADRA2C,CCDC59,CKAP4,CSF2RA,CSF2RB,CTSH,DMBT1,GATA6,LMCD1,NAPSA,P2RY2,PGA3,PGA4,PGA5,SFTA3,SFTPA1,SFTPA2,SFTPB,SFTPC,SFTPD,SLC34A1,SLC34A2,TTF1,ZDHHC2 |
| LipidOxidation | ABCB11,ABCC9,ABCD1,ABCD2,ABCD3,ABCD4,ACAA1,ACAA2,ACAD10,ACAD11,ACADL,ACADM,ACADS,ACADVL,ACAT1,ACOT8,ACOX1,ACOX2,ACOX3,ACOXL,ACSM1,ADH4,ADH5,ADH7,ADIPOQ,ADIPOR1,ADIPOR2,AKT1,AKT2,ALDH1L2,ALOX12,ALOX12B,ALOX15,ALOX15B,ALOX5,ALOXE3,AMACR,APOD,APPL2,AUH,BDH2,CRAT,CROT,CYGB,CYP4F2,CYP4F3,CYP4V2,DECR1,DECR2,DGAT2,ECH1,ECHDC1,ECHDC2,ECHS1,ECI1,ECI2,EHHADH,ETFA,ETFB,ETFBKMT,ETFDH,FABP3,FMO1,FMO2,FMO4,GCDH,GDF15,HACL1,HADH,HADHA,HADHB,HAO1,HAO2,HSD17B10,HSD17B4,ILVBL,IRS1,IRS2,IVD,KLHL25,LEP,LONP2,MAPK14,MCAT,MFSD2A,MLYCD,MTLN,PDK4,PEX13,PEX2,PEX5,PEX7,PHYH,PLA2G7,PLIN5,POR,PPARA,PPARD,PPARGC1A,SAMD1,SCP2,SESN2,SIRT4,SLC25A17,SLC27A2,SOX9,TWIST1,TYSND1 |
| AMPKmediatedFattyAcidOxidation | ACACB,ACSL1,CAB39,CAB39L,CPT1A,CPT1B,CPT2,PRKAA1,PRKAA2,PRKAB1,PRKAB2,PRKAG1,PRKAG2,PRKAG3,SLC25A20,STK11,STRADA,STRADB |
| EMT | ADAP1,ATP2C2,CLDN3,CLDN4,CLDN7,EHF,EPN3,ESRP1,ESRP2,GRHL1,GRHL2,IRF6,LLGL2,MARVELD2,MARVELD3,MYO5B,OVOL1,PRSS8,RAB25,S100A14,ST14,TJP3 |
| LipofibroblastSignature | AGO1,ALDH1A1,BMPER,CD59A,CDKN2C,CES1D,CRISPLD2,CCN1,DNAJB11,VPS26C,FHL1,ADGRG6,GSTM1,HSD11B1,IFI27L2A,IFITM1,IGFBP3,IGFBP6,INMT,ITGA8 |

Table S2: Curated gene sets per molecular feature for equine synovial screens

| OA Molecular Feature | Genes |
|---|---|
| Cartilage Matrix Degradation | *ADAMTS2,ADAMTS5,ADAMTS9,ADAMTSL4,ADAMTS6,ADAMTS14,ADAMTSL1,ADAMTSL3,ADAMTS7,ADAMTS12,ADAMTS4,ADAMTS16,ADAMTS17,ADAMTS10,ADAMTS1,ADAMTS19,ADAMTS3,ADAMTS15,ADAMTSL5,ADAMTS20,MMP12,MMP13,MMP23B,MMP25,ADAM17,ADSMTS11,MMP1,MMP14,MMP9,MMP11,MMP15,MMP16,MMP17,MMP19,MMP21,MMP28,ADAM9,MMP8,MMP3,MMP7,MMP10,MMP24,ADAM8,ADAM10,ADAM15,ADAMDEC1,ADAMTS8,ADAMTS18,ECE1* |
| Antioxidants | *ATP7A,SOD1,SOD2,SOD3,CAT,GPX3,GPX4,GPX7,GPX8,PRDX1,PRDX2,PRDX3,PRDX5,PRDX6,GSR,GSTP1,TXNRD2* |
| Cell Division | *CCNA1,CCNB1,CCNC,CCND1,CCND2,CCND3,CCNE1,CCNE2,CCNF,CCNG1,CCNH,CCNI,CCNJ,CCNJL,CCNK,CCNL1,CCNL2,CCNQ,CCNT1,CCNT2,CCNY,CCNYL1,CDKL3,CDKL5,CDK1,CDK2,CDK3,CDK4,CDK5,CDK6,CDK7,CDK8,CDK9,CDK10,CDK12,CDK13,CDK14,CDK16,CDK17,CDK19,CDK20,E2F1,E2F3,E2F4,E2F5* |
| Apoptosis | *APAF1,ATM,CASP1,CASP10,CASP2,CASP6,CRADD,PIDD1,TP53,TP63,AIFM2,BAX,BCL6,BID,BIRC5,BNIP3L,CHM,CREBBP,FAS,IGFBP3,NDRG1,PERP,PMAIP1,PPP1R13B,PRELID1,PRELID3A,RABGGTB,TMEM219,TP53BP2,TP53I3,TP53INP1,TRIAP1,ANXA6,ANXA4,ANXA11,ANXA13,TNFSF12,DISC1,CASP3,ANXA5,ANXA2,CASP9,BAD,BAK1,CASP8,CASP7,FADD,DAXX,BLK,DIABLO,GAS1,HIPK1,DDIT3,GADD34,BAK1,PPP1R15A,BCL2L11,GADD45G* |
| NLRP3 Inflammasome | *NLRP3,P2RX7,PANX1,PSTPIP1,PYCARD,SUGT1,TXNIP,ATAT1,CD36,DDX3X,DHX33,GBP5,NEK7,TLR6,NLRP1* |
| Reelin_Pathway | *FYN,LRP8,RELN,VLDLR,AKT1,GSK3B,MAP1B,MAPK8,MAPT,PAFAH1B1,PIK3CA,PIK3R1,LRPAP1,MAP1B,MAP2K7,MAP3K11,MAPK8,MAPT,NCK2,PAFAH1B1,RAP1A,RAPGEF1,SH3KBP1,CRK,DAB2IP,STAT5A* |
| Ion_Channels | *SCN2A,SCN8A,SCN9A,SCN4B,KCNMA1,ASIC3,CACNA2D1,CACNA1B,PIEZO1,COX6C,PCSK6,COMT* |
| Calcium_Binding | *ACTN1,ACTN2,ACTN3,ACTN4,ANXA1,ANXA10,ANXA11,ANXA13,ANXA2,ANXA3,ANXA4,ANXA5,ANXA6,ANXA7,ANXA8,ATP2A1,ATP2A2,ATP2A3,CALB1,CALB2,CALM1,CALM2,CALM3,CALML3,CALML4,CALML5,CALML6,CALR,CANX,CAPN1,CAPN2,CAPN3,CASQ1,CASQ2,CDH1,CDH11,CDH15,CDH2,CDH3,CDH5,DST,HPCAL1,HPCAL4,HRC,ITGA2B,ITGB3,NCS1,NOS1,NOS2,NOS3,OCM,PCDH1,PEF1,PLA2G4A,PLCD1,PLCD3,PLCD4,PRKCA,PRKCB,PRKCG,PVALB,RCVRN,RYR1,RYR2,S100A1,S100A10,S100A11,S100A12,S100A13,S100A14,S100A16,S100A2,S100A3,S100A4,S100A5,S100A6,S100A7,S100A8,S100A9,S100B,S100G,S100P,S100Z,SCGN,SORBS1,SORBS2,SRI,SYT1,SYT11,SYT2,SYT7,TGM1,TGM2,TGM3,TNNC1,TNNC2,VSNL1* |
| JAK STAT Activation | *IL6ST,JAK1,STAT3,CSNK2A1,STAT1,STAT6,JAK2,CEBPB,JAK3* |
| NFKB Activation | *RELA,NFKB1,NFKB2,RELB,CREL,IKBKG,IKBKB,CHUK* |
| IL1 Signaling | *CUL1,TAB2,MAP2K4,FBXW11,ALPK1,TOLLIP,PSMC5,N4BP1,RIPK2,NOD1,MAP3K8,MAP2K6,SKP1,IL1A,IL1R2,IL1R1,IL1B,SEM1,IRAK2,PELI2,NLRC5,TAB3,S100B,NLRX1,LRRC14,S100A12,IKBIP,BTRC,NOD2,NKIRAS2,TNIP2,MAP2K1,SAA1,PELI3,TRAF6,IRAK1,USP18,HMGB1,IL1RAP,PELI1,NKIRAS1,IRAK4,MAP3K3,AGER,MYD88* |
| IFNG Signaling | *CAMK2A,CAMK2B,CAMK2G,CIITA,FCGR1A,FCER1GP,GBP1,GBP2,GBP3,GBP7,H2-A,H2-B,H2-DMA,H2-DMB1,H2-DMB2,H2-EA,H2-EB1,H2-L,H2-M3,H2-OA,H2-Q7,H2-T24,HLA-A,HLA-DPA1,HLA-DPB1,HLA-DQB1,HLA-DRA,HLA-DRB1,HLA-DRB3,HLA-DRB4,HLA-DRB5,HLA-F,HLA-G,IFI30,IFI44,IFNGR1,IFNGR2,IFNG,IRF1,IRF3,IRF5,IRF7,IRF8,IRF9,MAPK1,MAPK3,MDA5,MICAL1,OAS1,OAS2,OAS3,PIM1,PLCG1,RHOB,TRIM10,TRIM12A,TRIM14,TRIM21,TRIM22,TRIM25,TRIM26,TRIM29,TRIM30A,TRIM34A,TRIM38,TRIM43,TRIM5,DRA,DQB,DRB,DQB* |
| TNFA Signaling | *AIM2,ATF2,CARD14,CARD16,CASP1,CASP3,CASP8,CD70,CCL2,EGFR,EPHA7R,F2R,F2RL1,FADD,FAS,GASC1,HSPA1A,HSPB1A,NLRC4,PPP2CB,PYDC1,RIPK1,RIPK3,SPHK1A,STK11,TAB1,TBK1,TIFA,TNFRSF14,TNFRSF1A,TNFRSF1B,TNFRSF12A,TNFRSF13B,TNFRSF13C,TNFRSF4,TNFRSF9,TNFSF12,TNFSF14,TNFSF15,TNFSF18,TRAF1,TRAF2,TRAF3,TRP53,TNFRSF10,TNFRSF12,TNFA* |
| Inflammatory mediators | *REACTOME_CHEMOKINE_RECEPTORS_BIND_CHEMOKINES,CXCL1,CXCL2,CXCL5,CXCL8,CXCL9,CXCL10,CXCL11,CXCL13,CXCL12,CXCL14,CCL2,CCL5,CCL7,CCL8,CCL11,CCL17,CCR2,CCR5,CCL19,CCL21,CCL20,CCL22,CCL17,IRF4,CCL3L1,CXCR4,CXCL1,CCL11,IL8,IL18,PLAU,LIF,CSF3,IL6,IL11,IL33,CSF1,CSF2,CSF3,NOS2,TNF,IL15* |
| Positive Regulation of ROS | *ADGRB1,CLCN3,CYBA,DUOXA1,DUOXA2,GRIN1,MIR24-1,PLCG2,RAB27A,SLC5A3,TLR4,ZNF205* |
| Negative Regulation of ROS | *ABCB7,ABCD1,ABCD2,ACP5,ATG5,BCR,BECN1,BNIP3,BRCA1,CFLAR,COA8,CRYAB,CTNS,FYN,G6PD,HDAC6,HIF1A,HK2,HP,INS,MIR181A2,MIR21,MIR590,MPV17L,MT3,PAGE4,PAX2,PINK1,PLIN5,PON3,PPARA,PRKN,RHOA,SIRT2,SIRT3,SIRT5,SLC18A2,TFAP2A,TIGAR,TRAP1,FOXO3* |
| NerveSensitizing Phenotype | *NGF,GDNF,BDNF,C3,CXCL1,CCL5,NTF3,ARTN,NT4,PTGS2,PTGS1,PANX1,KLK1,SPTLC1,SPTLC2,CCL20,LAMB3,TACR1,P2RX7,CSF1R,PIEZO2,PI16,TACR1,TAC1,BDKRB2* |
| ER Stress | *ERN1,EIF2AK3,ATF6,ATF4,XBP1,DDIT3,HSPA5,HSP90B1,CALR,CANX,PDIA3,PDIA4,PDIA6,ERN1,HSPA5,HSPA1A,HSPB1,HSPD1* |
| Hypoxia | HALLMARK_HYPOXIA with the following genes removed: *BGN,CCN1,CCN2,COL5A1,DCN,ENO2,GPC1,IGFBP3,JUN,LOX,PLAUR,SDC4,SERPINE1,TGFBI,TGM2,TNFAIP3,VEGFA* |
| Wnt Singaling | *FRAT1,FRAT2,RUVBL1,AKT1,EP300,SOX9,BTRC,CSNK2A1,CTNNBIP1,CSNK2A2,AXIN1,CSNK1E,DKK1,SFRP1,SFRP2,CSNK2B,TCF7,CUL1,LRP5,CXXC4,LRP6,PPP2CA,WNT6,WNT2,WNT3,WNT3A,CSNK1A1,CAV1,WNT7A,BCL9,TCF4,TCF3,SKP2,CCND2,HEY1,HEY2,GNAI1,NKD1,PPARD* |

| | |
|---|---|
| Fibrotic ECM | *AEBP1,CILP,COL1A1,COL1A2,COL3A1,COL5A1,COL5A2,COL6A1,COL6A3,COMP,FN1,FNDC1,POSTN,SPARC,THBS1,TNC* |
| Lipid Metabolism | REACTOME_FATTY_ACID_METABOLISM,HALLMARK_FATTY_ACID_METABOLISM,REACTOME_LIPID_DIGESTION_MOBILIZATION_AND_TRANSPORT |
| Oxidative Phosphorylation | GOBP_OXIDATIVE_PHOSPHORYLATION,REACTOME_RESPIRATORY_ELECTRON_TRANSPORT,REACTOME_MITOCHONDRIAL_RESPIRATION |
